# RNase-mediated reprogramming of *Yersinia* virulence

**DOI:** 10.1101/2024.01.11.575149

**Authors:** Ines Meyer, Marcel Volk, Ileana Salto, Theresa Moesser, Anne-Sophie Herbrüggen, Manfred Rohde, Michael Beckstette, Ann Kathrin Heroven, Petra Dersch

## Abstract

RNA degradation is an essential process that allows bacteria to regulate gene expression and has emerged as an important mechanism for controlling virulence. However, the individual contributions of RNases in this process are mostly unknown. Here, we report that of 11 tested potential RNases of the intestinal pathogen *Yersinia pseudotuberculosis*, two, the endoribonuclease RNase III and the exoribonuclease PNPase, repress the synthesis of the master virulence regulator LcrF. LcrF activates the expression of virulence plasmid genes encoding the type III secretion system (Ysc-T3SS) and its substrates (Yop proteins), that are employed to inhibit immune cell functions during infection. Loss of both RNases led to an increase in *lcrF* mRNA levels and stability. Our work indicates that PNPase exerts its influence via YopD, known to accelerate *lcrF* mRNA degradation. Loss of RNase III results in the downregulation of the CsrB and CsrC RNAs, leading to increased availability of active CsrA, which has previously been shown to enhance *lcrF* mRNA translation and stability. Other factors that influence the translation process and were found to be differentially expressed in the RNase III-deficient mutant could support this process.

Transcriptomic profiling further revealed that Ysc-T3SS-mediated Yop secretion leads to global reprogramming of the *Yersinia* transcriptome with a massive shift of the expression from chromosomal towards virulence plasmid-encoded genes. A similar extensive transcriptional reprogramming was also observed in the RNase III-deficient mutant under non-secretion conditions. This illustrates that RNase III enables immediate coordination of virulence traits, such as Ysc-T3SS/Yops, with other functions required for host-pathogen interactions and survival in the host.

**Author Summary:** Bacterial pathogens need to quickly adapt the expression of virulence- and fitness-relevant traits in response to host defenses. Pathogenic *Yersinia* species rapidly upregulate a type III secretion system (T3SS) to inject antiphagocytic and cell toxic effector proteins, named *Yersinia* outer proteins (Yops), into attacking immune cells. For this purpose, they display complex and resilient regulatory mechanisms. At the post-transcriptional level, this is mediated by different RNA-binding regulators including YopD and CsrA, while the fate of mRNAs is balanced by ribonucleases. Here, we demonstrate that out of 11 tested putative RNases of *Yersinia*, two major RNases, the endoribonuclease RNase III, and the exonuclease and degradosome component PNPase play a crucial role in the activation of the Ysc-T3SS/Yop machinery. We show that they promote the decay of the *lcrF* mRNA encoding the common transcriptional activator LcrF of the Ysc-T3SS/Yop components. PNPase seems to act through the control of the effector YopD, known to promote the decay of the *lcrF* transcript. In contrast, RNase III triggers processes that reduce *lcrF* mRNA translation and stability, and involve CsrA.

A transcriptome analysis further revealed that RNase III controls a series of events that include rapid and massive genetic reprogramming from mainly chromosomal-encoded genes to virulence-plasmid-encoded *ysc*-T3SS/*yop* genes. This control process does not only ensure immediate counter-measures during an immune attack, it also helps to overcome accompanying energetic and stress burdens and allows to rapidly readjust the genetic program after a successful defense.

## Introduction

Bacterial pathogens employ a large variety of regulators and integrate control mechanisms to tightly control the synthesis of crucial virulence factors on the transcriptional and post-transcriptional levels. In particular, RNA-mediated control processes provide pathogens with a highly efficient strategy to quickly respond to changes in their environment outside and inside their hosts [1][2][3]. Among these mechanisms are sensory and regulatory RNAs, which were shown to modulate virulence factor expression in response to external stimuli [1][2][3]. Other important players, which have been less addressed in this scenario, are ribonucleases (RNases). They play a critical role in RNA metabolism as they degrade RNA and are involved in the processing and quality control of transcripts [4]. RNases are enzymes that catalyze the cleavage of a phosphodiester bond within a double- or single-stranded RNA molecule (endonucleases) or from the 3’- or 5’-end (exonucleases) [5][6]. While some RNases are highly conserved across different species (e.g. RNase E, PNPase), some are only present in certain bacteria, e.g. RNase III in Gram-negative bacteria. The susceptibility of a particular RNA to an RNase thereby depends on multiple factors: (i) intrinsic RNA properties, e.g. RNA secondary structure, (ii) rate of translation, (iii) RNA-binding proteins, and (iv) sRNAs [4][7][8][9][10].

More recently, certain RNases have also been found to be crucial regulators of virulence genes. Among them are specific exonucleases, i.e. PNPase, and endo– ribonucleases such as RNase Y in Gram-positive bacteria (*Staphylococcus aureus, Streptococcus pyogenes,* and *Clostridium perfringens*) or RNase III in Gram-negative bacteria (*Escherichia coli*, *Salmonella enterica* serovar Typhimurium) [7][8][9][11]. Roughly 15-20 different RNases are encoded in a single bacterium, and the functions of some of the enzymes are coregulated and can be complemented and/or compensated to a certain degree by others [12][13][14]. Considering the essential role of RNases in RNA metabolism, it is expected that the expression and the activity of RNases are tightly regulated [5][15][16]. Bulk RNA degradation was found to be dependent on a membrane-bound, multiprotein complex, the degradesome, including RNases, such as RNase E and the polynucleotide phosphorylase (PNPase), and auxiliary proteins, which assist and/or regulate the degradation process. The composition of the degradosome varies between different bacteria and can change in response to environmental conditions [7][8][17]. Both degradosome components RNase E and PNPase are involved in the control of *Yersinia* virulence [18][19][20][21].

*Yersiniae* are Gram-negative bacteria of which *Y. pestis*, the causative agent of plague, and its enteric relatives *Y. enterocolitica* and *Y. pseudotuberculosis* are human pathogens. *Y. pseudotuberculosis,* the most closely related, direct ancestor of *Y. pestis* is an intestinal pathogen [22]. It causes a variety of gut-associated diseases, such as watery diarrhea, enteritis, and mesenteric lymphadenitis, called Yersiniosis. Upon oral uptake, the bacteria enter and transfer through the intestinal epithelial layer into the underlying lymphatic tissues (Peyer’s patches) where they rapidly proliferate extra– cellularly [23][24][25]. During this infection phase, the *yersiniae* induce the expression and formation of the Ysc type III secretion system (Ysc-T3SS) encoded on the 70 kb virulence plasmid pYV/pIB1 [26][27]. The Ysc-T3SS, a syringe-like machinery called the injectisome injects 7-8 Yop effector proteins into professional phagocytes to protect the bacteria from phagocytosis, manipulate inflammatory processes, and induce cell death [28][29]. Expression of the Ysc-T3SS/Yop machinery is tightly regulated in response to temperature and host cell contact by a complex network of regulatory factors [22][26][18][30]. The majority of the *ysc*/*yop* genes is induced by the AraC-type transcriptional activator LcrF [31]. Synthesis of LcrF is thermally regulated on the transcriptional and post-transcriptional levels. This is mediated by the thermo-sensitive transcriptional repressor YmoA and a *cis*-acting RNA thermometer element including the ribosomal binding site of *lcrF* [32][33]. Synthesis of the Ysc-T3SS/Yop apparatus is also tightly linked to the activity of the secretion machinery. Pettersson *et al*. demonstrated that expression of the injectisome components and the Yops are strongly induced upon host cell contact [30]. Cell contact-mediated induction can be mimicked by depletion of Ca^2+^, a phenomenon known as the low calcium response. This converts the T3SS apparatus into the secretion state and enables a strong, synchronized activation of the *ysc*-T3SS and *yop* genes [34][35]. In a previous study, we demonstrated that one of the secreted Yop proteins, YopD, plays a pivotal role as a host cell contact sensing device and negative regulator of Ysc-T3SS/Yop synthesis in the absence of inducing cues [18]. YopD was found to coopt important global riboregulators to control the Ysc-T3SS/Yop master regulator LcrF. Secretion of the YopD translocator upon cell contact increases the ratio of the post-transcriptional RNA-binding regulator CsrA to its antagonistic small RNAs CsrB and CsrC. YopD secretion also reduces the levels of the degradosome RNases PNPase and RNase E [18]. This indicated that riboregulators and RNA-mediated control processes play a pivotal role in the coordinated control of *Yersinia* virulence traits.

In this study, we extended our analysis and found that in particular RNase III and PNPase reduce the stability of the *lcrF* mRNA. PNPase exerts its main influence on Ysc-T3SS/Yop synthesis through the RNA-regulatory function of YopD [18], whereas RNase III influences translation and stability of *lcrF* mRNAs via (i) CsrA-antagonizing Csr RNAs, and (ii) likely by the control of ribosomal and translation-related factors.

## Results

### Identification of RNases implicated in type III secretion of *Y. pseudotuberculosis*

In our attempt to identify RNases that influence virulence traits of *Y. pseudotuberculosis*, in particular the expression of the T3SS machinery, we constructed in-frame deletions in genes of identified putative RNases (*rnb*, RNase II; *rnc*, RNase III, *rnd*, RNase D; *rng*, RNase G; *rnhA*, RNase HI; *rph*, RNase PH; *yoaB*, L-PSP(YoaB); *tcdF,* TdcF; *tcdF-1,* TdcF-1; *tcdF-2,* TdcF-2; and *pnp* PNPase), except RNase E as its loss is lethal [19]. The chosen RNases differ in their target specificity, cleavage mechanism, and role in bacterial physiology [9][10], and previous global expression analysis by RNA-seq showed that these RNases are all controlled in response to virulence-relevant parameters such as temperature and/or nutrient availability [36][37].

A first phenotypic characterization of the RNase mutants demonstrated that the PNPase and RNase III-deficient strains (YP139 (Δ*pnp*), YP356 (Δ*rnc*)) exhibited a small growth delay at 25°C. However, in contrast to the Δ*pnp* mutant, the growth defect of the *rnc* mutant was much stronger at 37°C. Complementation by the respective RNase gene could restore the phenotype of the wildtype (Fig. **1A-B**). A similar growth arrest has been observed upon a constitutive expression of the Ysc-T3SS of *Yersini*a [38][39][40]. We further tested growth at 37°C under Ca^2+^-limiting conditions, as a substitute for host cell contact, to induce Ysc-T3SS/Yop expression and Yop secretion (secretion conditions). In agreement with previous results [38][39][40], induction of Yop secretion is accompanied by a strong growth reduction of the wildtype which was more severe in both RNase-deficient mutants (Fig. **1A**). All other tested RNases did not influence bacterial growth (Supplementary Fig. **S1A**-**B**).

**Figure 1:**
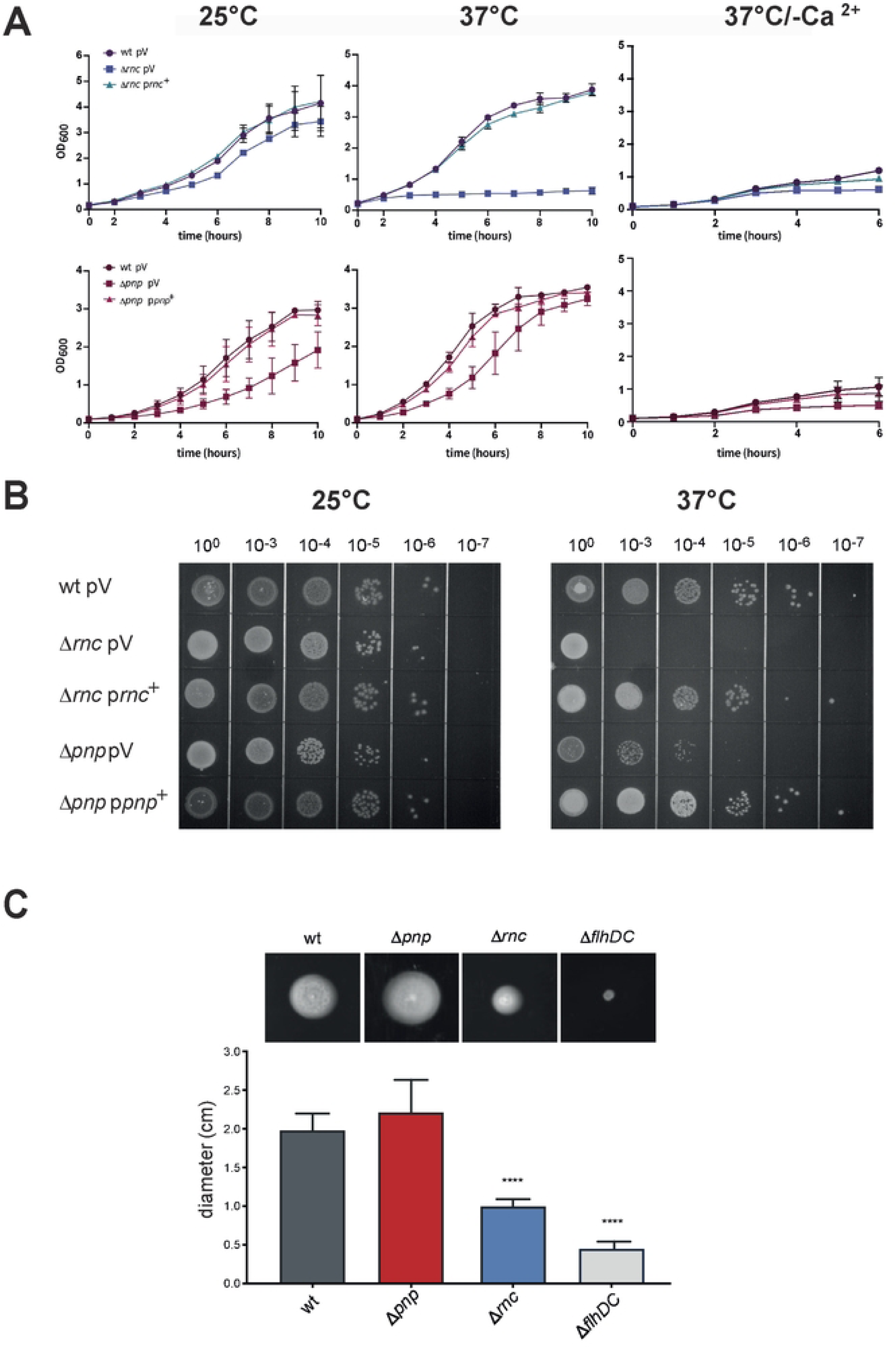
Loss of RNases PNPase and RNase III affects growth and motility of *Y. pseudotuberculosis*. (**A**) Overnight cultures of *Y. pseudotuberculosis* YPIII and the isogenic RNase mutants YP139 (Δ*pnp*) and YP356 (Δ*rnc*) harboring the empty vector (pV) or the complementation plasmids (p*rnc*^+^ or p*pnp*^+^) were diluted to an OD_600_ of 0,2 in LB and growth at 25°C, 37°C, and 37°C/-Ca^2+^ was followed by measurement of OD_600_. Data represent the mean ± SD from experiments done in triplicates. (**B**) One representative example of three independent spotting assays of serial dilutions of strains, used in A, on LB agar plates incubated at 25°C and 37°C overnight. (**C**) An equal amount of *Y. pseudo-tuberculosis* wildtype YPIII and the RNase mutants YP139 (Δ*pnp*) and YP356 (Δ*rnc*) was spotted onto tryptone swarm semi-soft agar plates and incubated at 25°C. A Δ*flhD*C mutant was used as negative control (upper panel). The mean ± SD of the diameters of the bacterial colonies is shown from experiments done in triplicates (lower panel). Statistical significance was determined using Student’s t-test. Asterisks indicate results significantly different from the wildtype; ****P<0.0001.

Moreover, motility, which is promoted by the flagellar T3SS, was found to be significantly reduced for the Δ*rnc* mutant grown on semi-solid agar plates at 25°C but not as drastic as the non-motile *flhDC* mutant missing the main activator of the flagellar synthesis genes (Fig. **1C** [41]). In contrast, a fuzzier, but not significantly larger concentric swarm ring was observed for the Δ*pnp* mutant strain (Fig. **1C**), whereas the motility of all other RNase mutant strains was not significantly affected (Supplementary Fig. **S1C**). Scanning electron microscopy revealed that wildtype and PNPase-deficient bacteria contained two-three flagella of the bacteria grown at 25°C, whereas the Δ*rnc* mutant largely lacked flagella at 25°C (Fig. **2A**). This indicated that motility loss of the Δ*rnc* mutant did result from the reduction of flagella and not of an inability to use flagella (i.e. a Mot-negative phenotype). In agreement with previous studies demonstrating a down-regulation of the transcription of flagellar genes at body temperature [42][43], flagella are absent at 37°C in all tested strains (Fig. **2A**). However, in contrast to the wildtype and the Δ*pnp* mutant, additional surface-exposed structures resembling needles of T3S injectisomes [44] were detected at 37°C with scanning and negative staining electron microscopy, which are also seen with the wildtype under secretion conditions (37°C/-Ca^2+^) (Fig. **2A-B**). This indicated that the production of the Ysc-T3SS/Yop machinery is induced at 37°C in the Δ*rnc* mutant, conditions under which the production and assembly of the apparatus are normally repressed [35][45].

**Figure 2:**
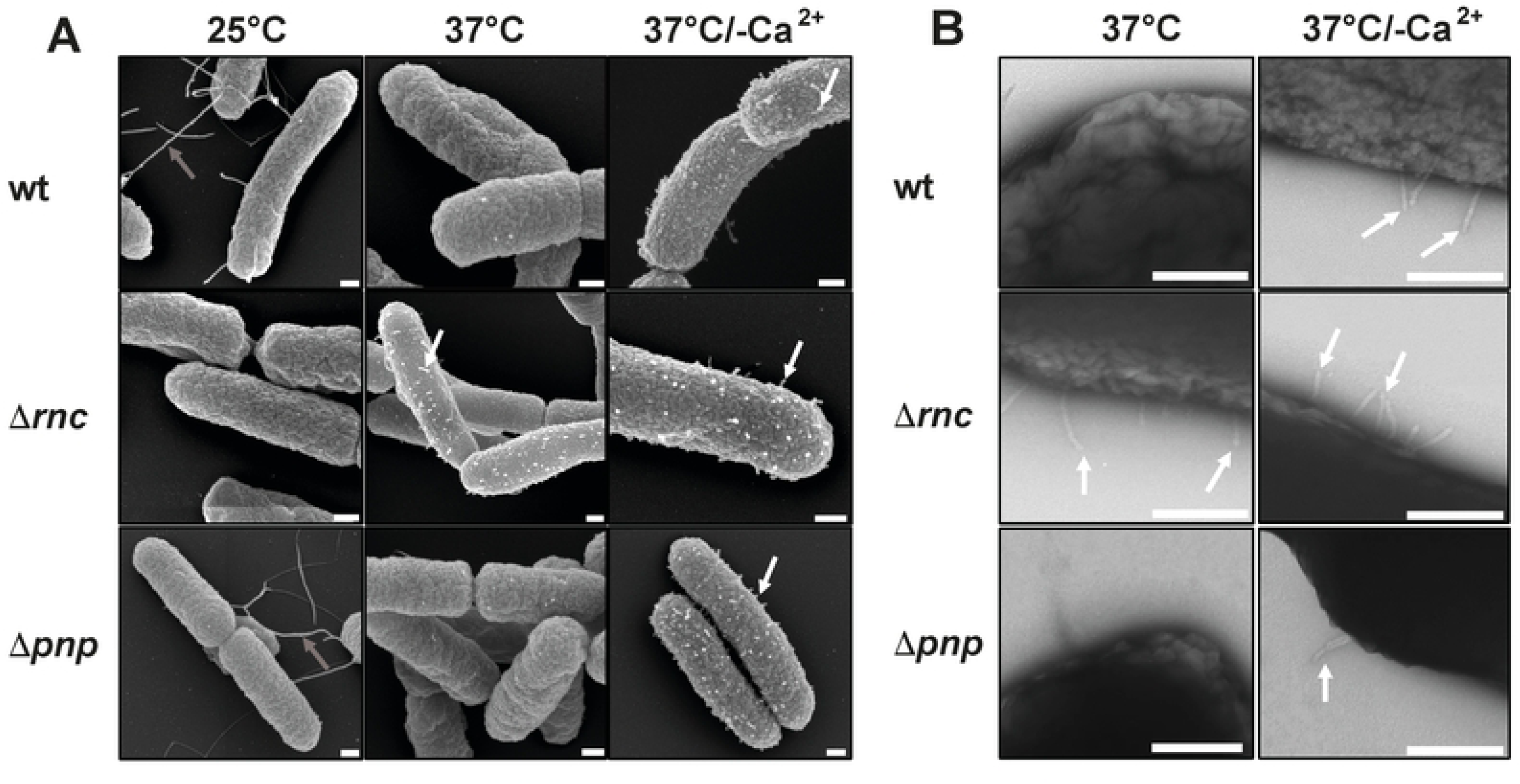
RNase III-dependent synthesis of flagella and T3S-like structures. Scanning (**A**) and transmission (**B**) microscopy of *Y. pseudotuberculosis* strains YPIII (wt), YP139 (**Δ***pnp*), and YP356 (**Δ***rnc*) grown at 25°C, 37°C and 37°C/-Ca^2+^. Flagella are indicated by grey and T3S injectisome-like structures by white arrows, respectively. Bars indicate 200 nm.

### Analysis of the influence of RNase III and PNPase on Yop secretion and expression

In order to test and characterize the influence of the PNPase and RNase III on the Ysc-T3SS/Yop machinery, *Y. pseudotuberculosis* wildtype strain YPIII and the isogenic Δ*pnp* and Δ*rnc* mutants were cultivated at 25°C (environmental, non-expression conditions), 37°C (host, non-secretion conditions) and 37°C/Ca^2+^-limiting conditions (mimicking host cell contact/secretion conditions) to trigger expression and activity of the Ysc-T3SS/Yop machinery [35][45]. A mutant strain deficient of the T3SS component YscS was used as a negative control as the encoded protein is known to be essential for Yop secretion [46]. Yop secretion by the wildtype was only detectable at 37°C/-Ca^2+^ (secretion conditions), but not at 25°C and 37°C (Fig. **3A**). In contrast, slightly higher amounts of Yops were already secreted by the Δ*pnp* mutant under non-secretion conditions (37°C), and a significantly higher amount of secreted Yop proteins was detectable under identical conditions in the absence of RNase III (Fig. **3A**). This did not result from bacterial cell lysis (Supplementary Fig. **S2A**) and the derepressed effect of the Δ*rnc* mutant could be complemented by an *rnc*-positive plasmid (Supplementary Fig. **S2B**). The observed Yop secretion pattern is in full agreement with the detection of numerous T3S injectisome-like structures on the surfaces of the bacteria (Fig. **2**). This might also explain the growth reduction observed at 37°C for the Δ*rnc* mutant (Fig. **1A, B**), the phenotype which is characteristic for Yop secretion conditions (37°C/-Ca^2+^) [38][39][40].

**Figure 3:**
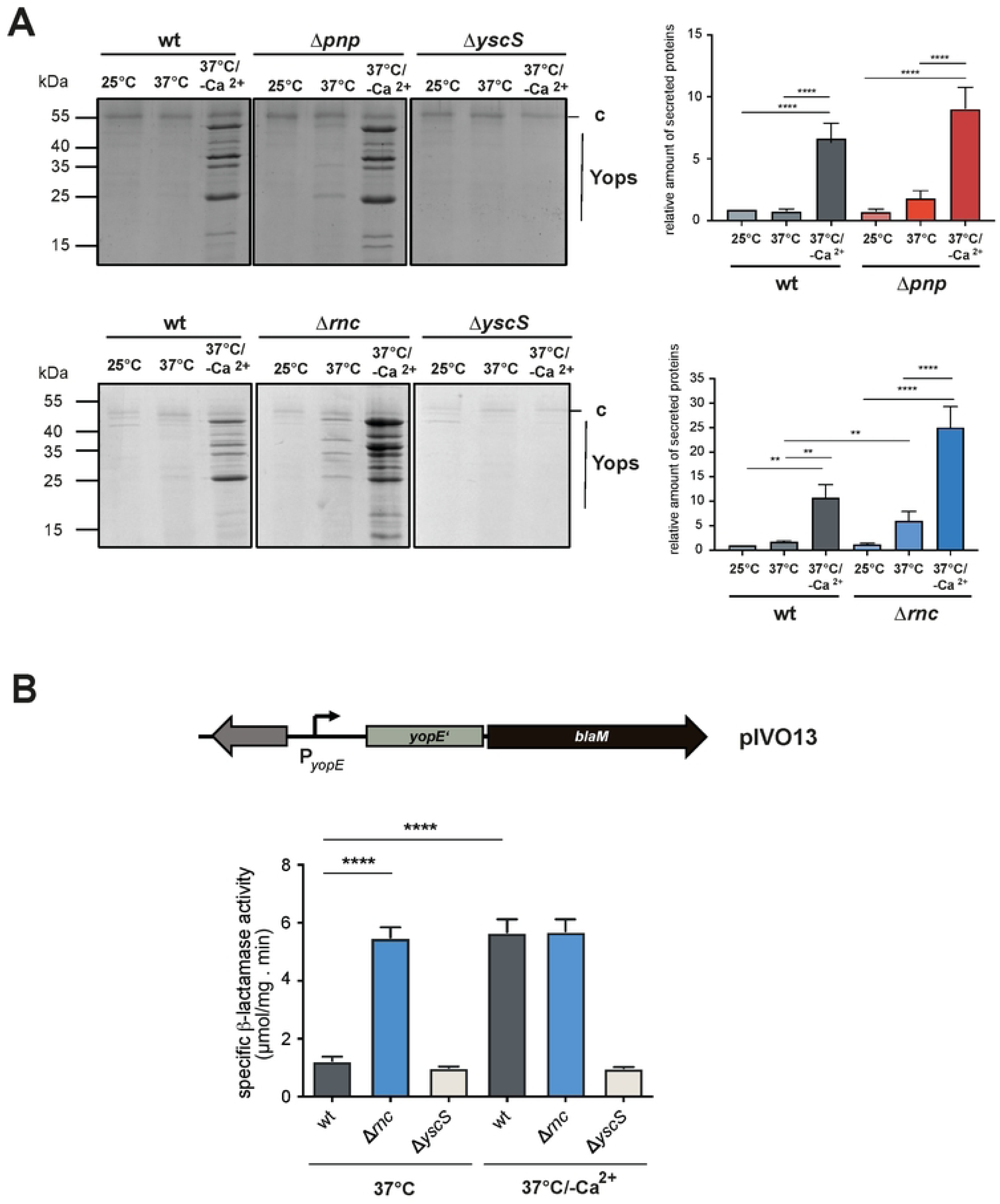
Influence of PNPase and RNase III on Yop protein secretion. (**A**) *Y. pseudotuberculosis* strains YPIII (wt), YP139 (Δ*pnp*), and YP356 (Δ*rnc*) were grown at 25°C, 37°C, and 37°C/-Ca^2+^; the secreted proteins in the supernatant of the cultures were precipitated with TCA and separated on SDS gels (left panels). The Yop secretion-deficient mutant YP101 (Δ*yscS*) was used as negative control. Secreted Yop proteins were quantified using ImageJ. Data represent the mean ± SD from three independent biological replicates relative to the amounts of Yops secreted by the wildtype at 25°C (right panel). Significant differences were determined using the one-way Anova test and are indicated by asterisks (*P <0.05, **P = 0.01). (**B**) Secretion of a plasmid-encoded YopE-beta-lactamase (BlaM) fusion protein (illustrated in the upper panel) expressed in *Y. pseudotuberculosis* strains YPIII (wt), and YP356 (Δ*rnc*) grown at 37°C and 37°C/-Ca^2+^ for 4 h was determined. Data represent the mean ± SD from three independent biological replicates (lower panel). Significant differences were determined using the Student’s t-test and are indicated by asterisks (****P<0.0001).

To confirm the influence of RNase III on the secretion of a specific Yop protein, we quantified the secretion of an introduced, plasmid-encoded YopE-β-lactamase fusion protein (YopE-BlaM) by nitrocefin assays after a shift from 25°C to 37°C +/- Ca^2+^ (Fig. **3B**). In agreement with our previous results, secretion of the YopE-BlaM fusion protein was significantly increased in the Δ*rnc* mutant at 37°C, but not in the wildtype, whereas similar high amounts of secreted YopE-BlaM were detectable with both wildtype and Δ*rnc* mutant under secretion conditions (37°C/- Ca^2+^).

To test, whether the intracellular amount or solely secretion of Yop proteins is affected, Yop protein levels in whole cell extracts of *Y. pseudotuberculosis* YPIII (wt), and the Δ*pnp* and Δ*rnc* RNase mutant strains grown under secretion (37°C/-Ca^2+^) or non-secretion conditions (37°C) were analyzed by Western blotting using an anti-all Yop antibody (Fig. **4**). This revealed that considerably higher amounts of the Yops are synthesized in the Δ*pnp* and the Δ*rnc* mutant at 37°C/non-secretion conditions. The overall Yop protein levels of both mutants are almost identical under secretion and non-secretion conditions and comparable to the Yop levels of the wildtype under secretion conditions (37°C/-Ca^2+^); except for YopE. This effector seems to be considerably reduced in the absence of RNase III compared to the wildtype (Fig. **4B-C**). Notably, similar to the other Yop proteins, the expression of the *yopE-blaM* reporter in the Δ*rnc* strain was strongly increased at 37°C (Fig. **3C**), whereas the YopE protein was barely detectable in the same strain (Fig. **4C**). This indicated that not *yopE* expression but rather post-transcriptional steps of YopE synthesis/stability are affected in the absence of RNase III.

**Figure 4:**
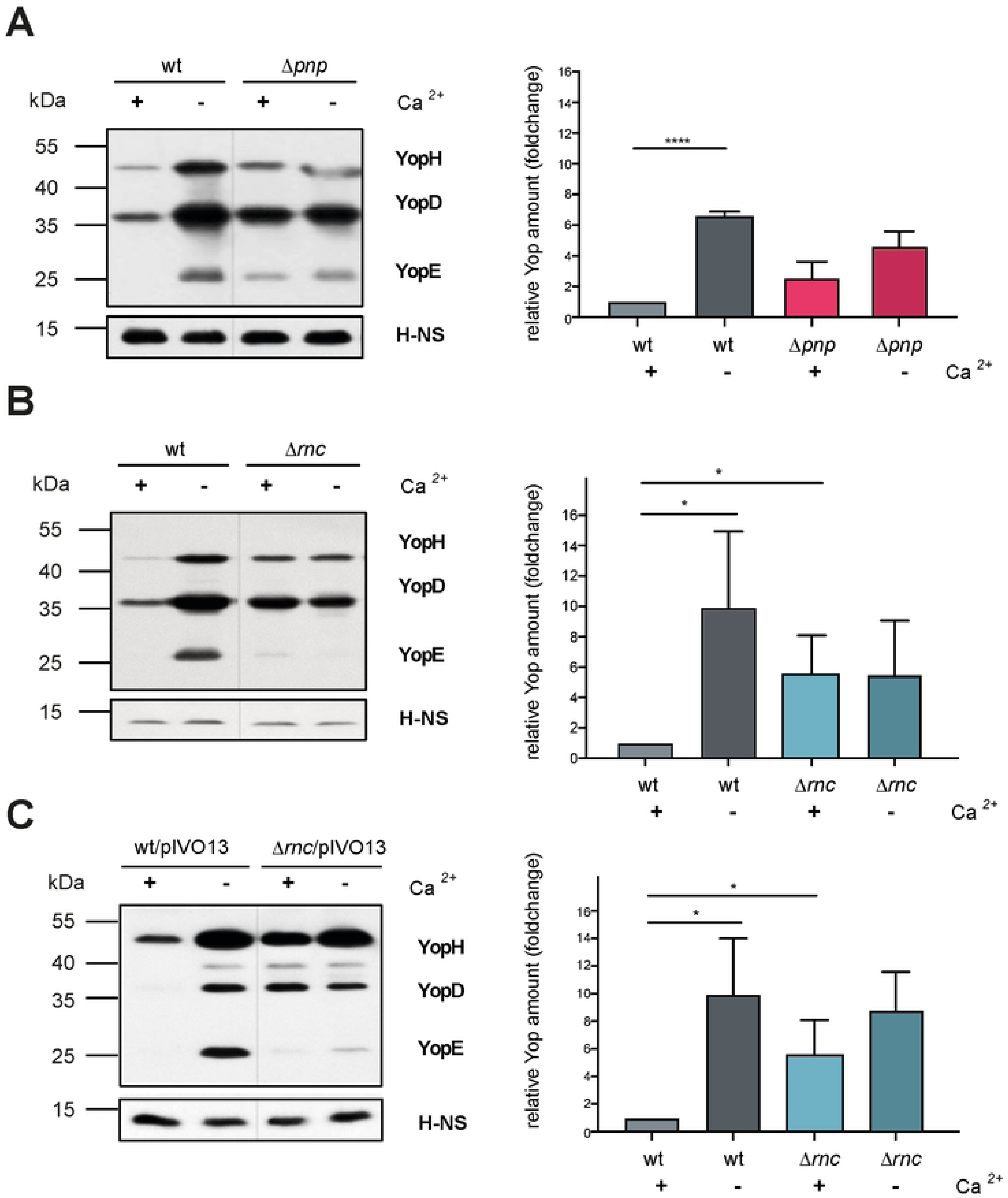
Yop synthesis is altered in PNPase- and RNase III-deficient strains. *Y. pseudotuberculosis* strains YPIII (wt), YP139 (Δ*pnp*), and YP356 (Δ*rnc*) without (**A**,**B**) or with a plasmid-encoded YopE-beta-lactamse (BlaM) fusion protein (**C**) were grown at 37°C (+) and 37°C/-Ca^2+^ (-); whole cell extracts were prepared and separated on SDS gels (left panels). Synthesized Yop proteins in the YPIII (wt) (**A-C**), YP139 (Δ*pnp*) (**A)**, YP356 (Δ*rnc*) (**B**), and (**C**) YP356 (Δ*rnc*) pIVO13 (*yopE*-*blaM*) were detected by Western blotting using a multi-Yop antiserum (left panel) and were quantified by ImageJ (right panel). An antiserum against H-NS was used for loading control. Data represent the mean ± SD from three independent biological replicates and relative amounts are documented with respect to the wildtype grown at 37°C. Significant differences were determined using Student’s t-test and indicated by asterisks (*P <0.05; ****P<0.0001).

To elucidate how the loss of PNPase and RNase III triggers the synthesis of the Ysc-T3SS/Yop components under non-inducing conditions, we next addressed the impact of the *pnp* and *rnc* knock-out mutations on known Ysc-T3SS/Yop control mechanisms and regulators.

### PNPase and RNase III influence on Ysc-T3SS/Yops does not occur through an increase of the pYV copy number

Synthesis of the Ysc-T3SS/Yop injectisome components is controlled on different levels, including the regulation of the copy number of the virulence plasmid pYV (also named pIB1) in response to temperature and host cell contact/secretion [47]. As the copy number of plasmids is subjected to complex regulation involving RNases [48], we employed qPCR to determine whether a knockout of the *rnc* or the *pnp* gene influences the pYV copy number. However, similar to the wildtype, we found only a subtle increase of the copy number from 1 to 2 copies in the mutants upon a shift from 25°C to 37°C (Supplementary Figure **S3**). Accordingly, increased Yop expression and synthesis at 37°C (Fig. **3**,**4**) cannot be explained by an increase in the pYV copy number and must be based on the deregulation of other control factors.

### RNase III and PNPase mainly influence Yop protein expression through the control of LcrF

Expression of the *yop* and the Ysc-T3S component genes is mainly activated by the AraC-type transcriptional activator LcrF in response to temperature and host cell contact [18][31][32]. Therefore, we tested whether the absence of RNase III and PNPase influenced LcrF synthesis. We found that the amount of the LcrF protein was considerably increased at 37°C/non-inducing conditions in both RNase mutant strains (Fig. **5A-C**). The overall influence of the loss of RNase III was more pronounced compared to the loss of PNPase (Fig. **5A-B**), and could be fully complemented by the *rnc*-encoding plasmid (Fig. **5C**). Interestingly, high LcrF levels in the Δ*rnc* strain at 37°C (no secretion) and 37°C/-Ca^2+^ (secretion) were comparable, demonstrating that LcrF synthesis is independent of the secretion control process in the absence of RNase III (Fig. **5B-C**).

**Figure 5:**
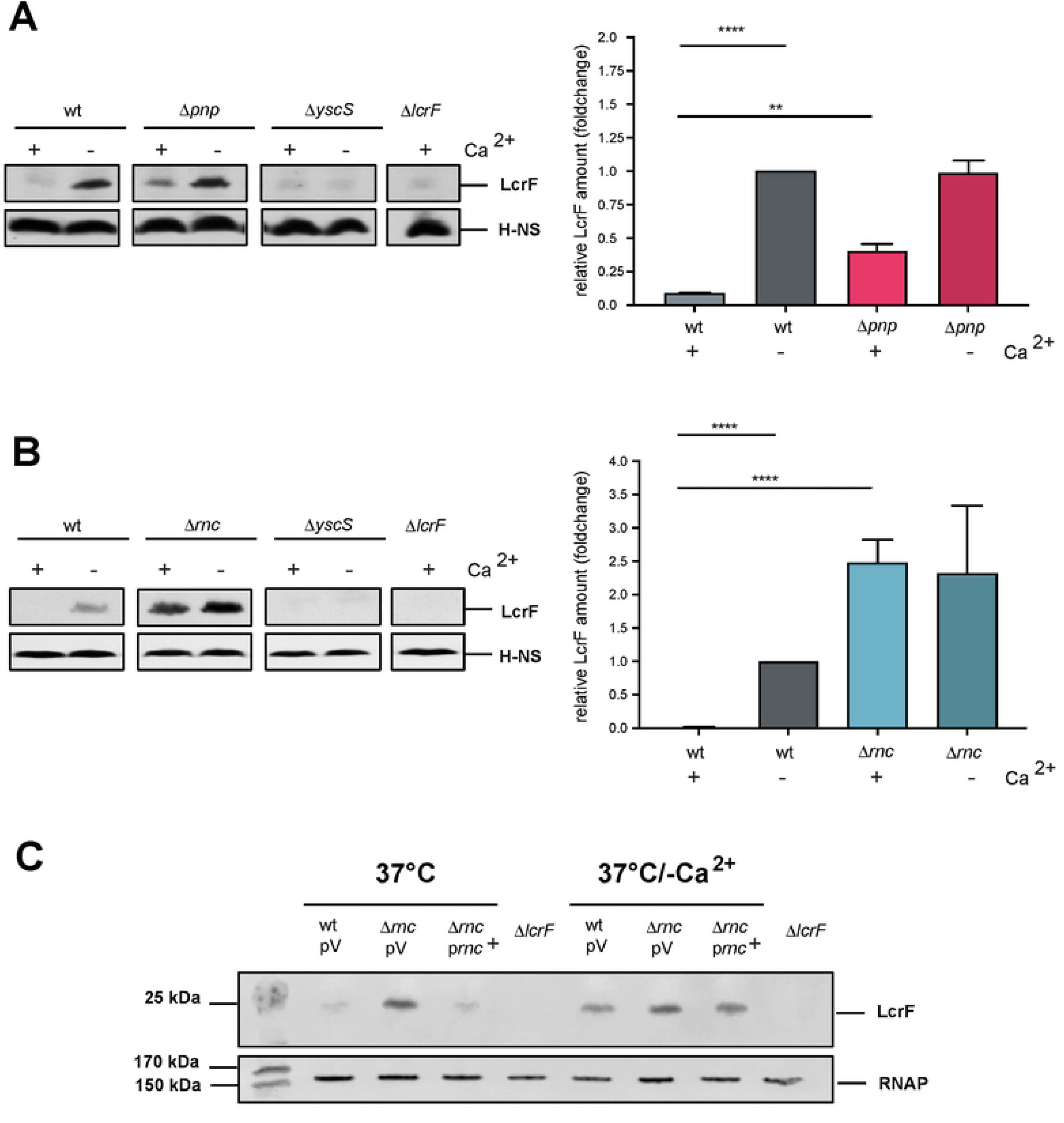
Influence of RNase III and PNPase on the synthesis of LcrF. *Y. pseudotuberculosis* strains YPIII (wt), YP139 (Δ*pnp*) (**A**), and YP356 (Δ*rnc*) (**B**) were grown at 37°C (+) and 37°C/-Ca^2+^ (-). The secretion-deficient mutant YP101 (Δ*yscS*) and the LcrF-deficient mutant strain YP179 (Δ*lcrF*) were used as a negative control. Whole-cell extracts were prepared and separated by SDS-PAGE. The transcriptional activator LcrF was detected by Western blotting using an LcrF-specific polyclonal anti-serum. An antiserum against H-NS was used for loading control (left panel). Data were quantified by ImageJ and represent the mean ± SD from three independent biological replicates (right panel). Significant differences were determined using Student’s t-test and indicated by asterisks (*P <0.05; ***P<0.001). (**C**) *Y. pseudotuberculosis* strains YPIII (wt), YP139 (Δ*rnc*), and YP356 (Δ*rnc*) pIV20 (p*rnc*^+^) were grown at 37°C (+) and 37°C/-Ca^2+^ (-); the LcrF-deficient mutant strain YP179 was used as a negative control. Whole-cell extracts were prepared and separated on SDS gels. Synthesized LcrF was detected by Western blotting using a polyclonal LcrF antiserum. An antiserum against RNA polymerase (RNAP) was used for loading control.

### RNase III and PNPase influence *lcrF* transcript levels

To unravel the molecular mechanism by which the RNases control LcrF levels, we first investigated the *lcrF* transcript level in the Δ*rnc* and Δ*pnp* mutant compared to wildtype at 37°C. In agreement with previous studies [18][32], we observed that the *lcrF* transcript is generally very unstable in the wildtype as visualized by a diffuse pattern of different-length transcripts hybridizing with the *lcrF* probe on Northern blots (Fig. **6A**). Although this general pattern did not change in the absence of both RNases a significantly higher amount of *lcrF* transcripts was observed in the absence of RNase III at 37°C (Fig. **6A**), and this effect could be fully complemented by a plasmid-encoded copy of the *rnc* gene (Fig. **6B**). Loss of PNPase caused a smaller upregulation of *lcrF* mRNAs under these conditions, whereas no influence of the RNases was observed under secretion conditions at 37°C/-Ca^2+^ (Fig. **6A**).

**Figure 6:**
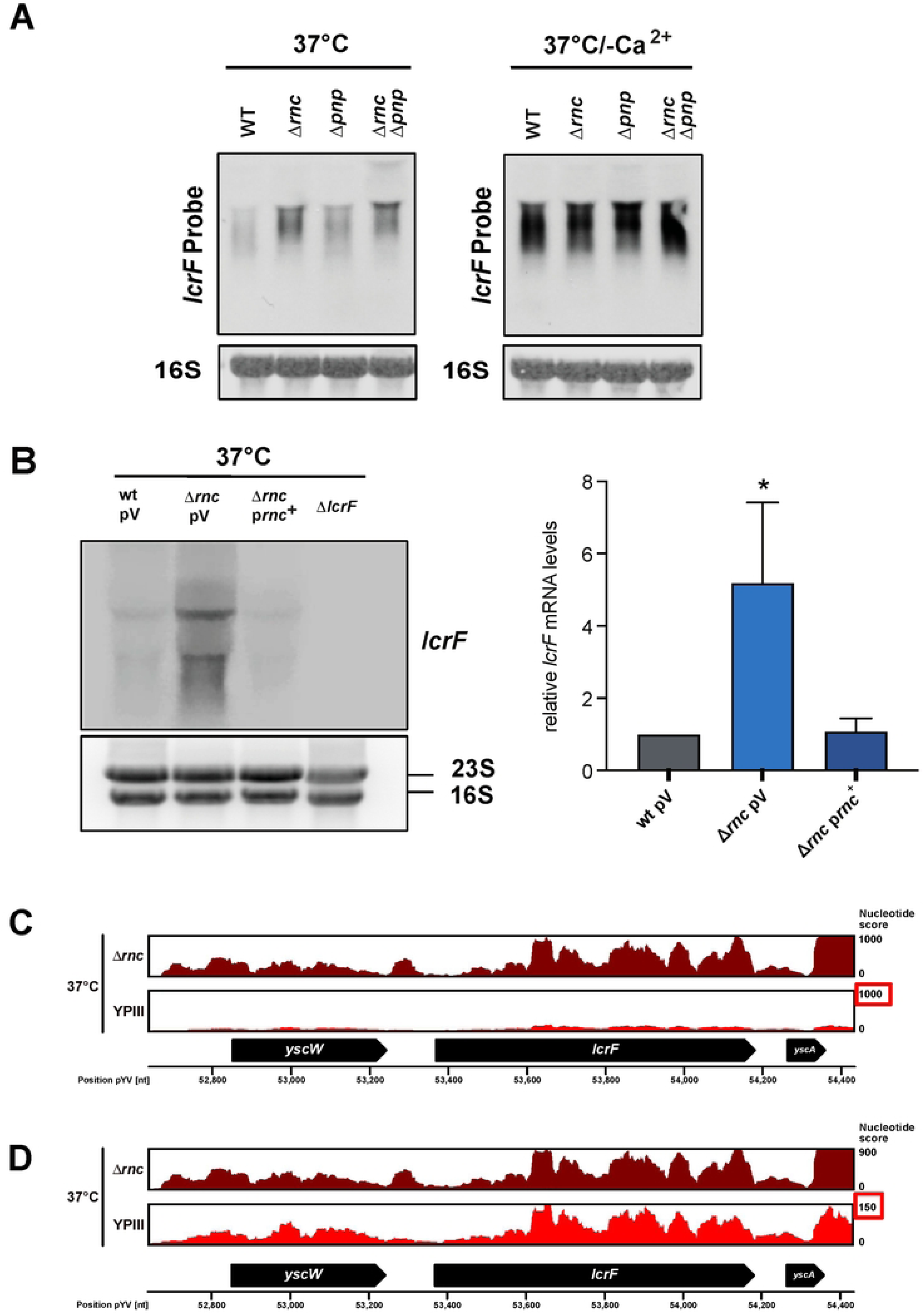
Influence of RNase III and PNPase on *lcrF* transcript levels. (**A**) *Y. pseudotuberculosis* strains YPIII (wt), YP139 (Δ*pnp*), YP356 (Δ*rnc*), and YP375 (Δ*rnc,* Δ*pnp*) were grown at 37°C and 37°C/-Ca^2+^. Total RNA of the samples was prepared and the *lcrF* transcript was detected by Northern blotting; 16S rRNA was used as loading control. (**B**) *Y. pseudotuberculosis* strains YPIII (wt), YP139 (Δ*pnp*), YP356 (Δ*rnc*), and YP356 (Δ*rnc*) pIVO20 (p*rnc*^+^) and the Δ*lcrF* strain (YP179) were grown at 37°C. Total RNA of the samples was prepared and the *lcrF* transcript was detected by Northern blotting (left panel) and quantified by ImageJ (right panel). Data represent the mean ± SD from three independent biological replicates. Significant differences were determined using Student’s t-test and indicated by asterisks (*P<0.05). (**C**) RNA-Seq coverage tracks of the *yscW-lcrF* locus in *Y. pseudotuberculosis* YPIII (wt) and YP356 (Δ*rnc*) grown at 37°C. RNA-Seq reads were mapped to the pYV virulence plasmid (NC_006153.2) and analyzed using a DESeq2 pipeline. The mapping visualizes the nucleotide score of the combined biological replicates for the two strains with an equal score for both. (**D**) RNA-Seq coverage tracks of the *yscW-lcrF* locus of the YPIII (wt) and YP356 (Δ*rnc*) were adjusted to nucleotide scores of 900 and 150, respectively to compare the read pattern of both strains.

In a parallel experiment, we also performed a comparative transcription profile analysis of the wildtype and the Δ*rnc* mutant grown at 37°C by RNA-sequencing (for more detail see also below). In agreement with the results of the Northern blots, we could detect a massive increase of reads mapping to *lcrF* in the RNA-seq data set (Fig. **6C**; Supplementary Dataset **S4**).

Upregulation of *lcrF* transcript levels in the Δ*rnc* and Δ*pnp* mutant could have occurred by the direct influence of the RNases on the stability of the *lcrF* mRNA or by an indirect effect on the activation of *lcrF* transcription. We first tested whether the overall increase of *lcrF* transcript levels in the absence of RNase III and PNPase is indirect and caused by an increase of *lcrF* transcription. For this purpose, we first used two reporter constructs, which both harbor the promoter region of the *yscW-lcrF* operon from -311 to +8 fused to *phoA* (transcriptional fusion, pMV53) and from -304 to +281 (including the entire 5’ untranslated region) relative to the transcriptional start site fused to *lacZ* (translational fusion, pKB35) to compare the activity of the promoter at 37°C. However, no increase in promoter P*_yscW-lcrF_* activity was detectable with both reporters in the Δ*rnc* and the Δ*pnp* mutant, respectively (Fig. **7A-B**). This demonstrated that an influence of the RNases on the initiation of *lcrF* transcription can be excluded.

**Figure 7:**
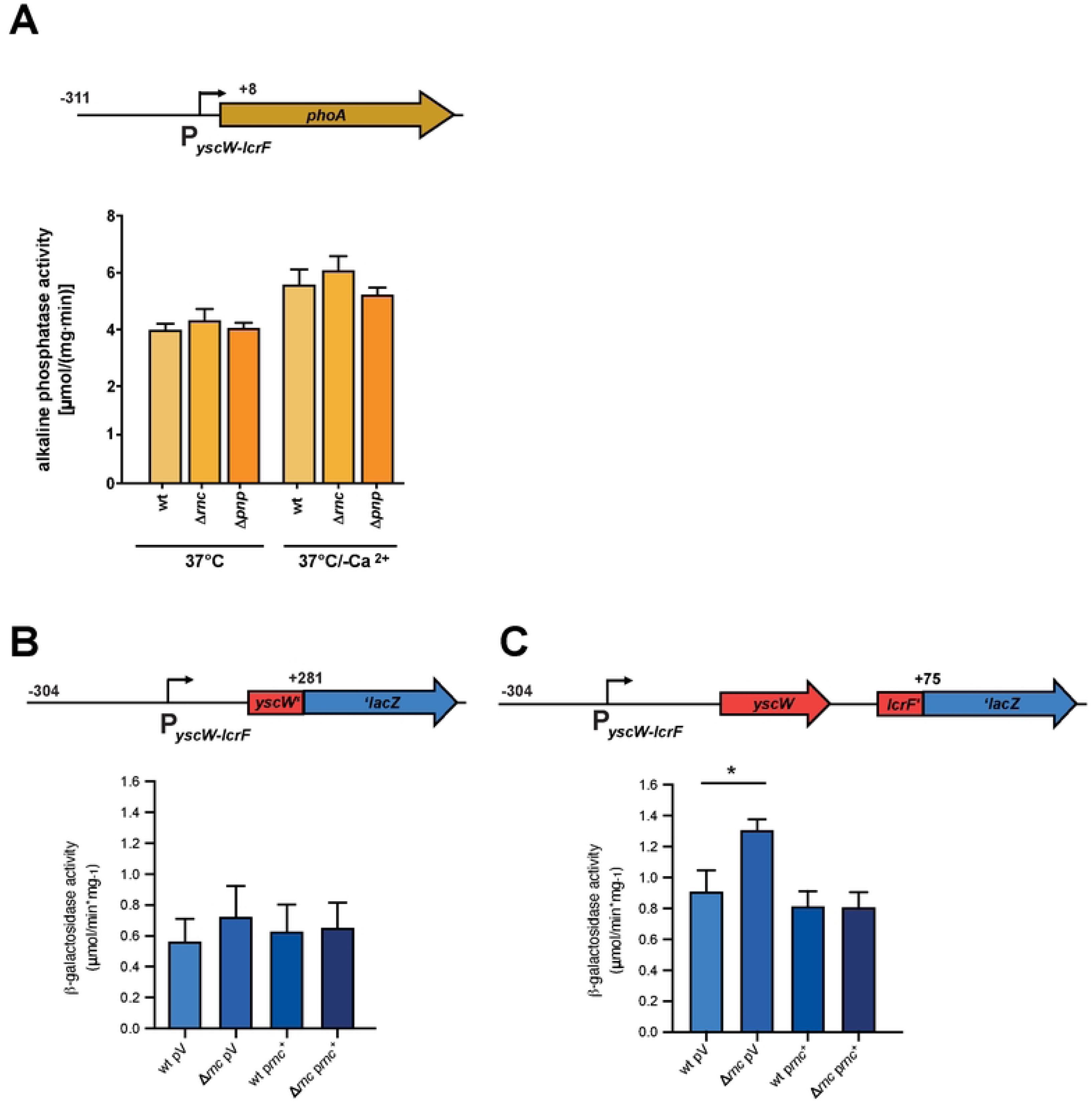
Influence of RNase III on *lcrF* transcription. (**A**) Plasmid pMV53 encoding the *yscW-phoA* transcriptional fusion and (**B**) plasmids pKB35 and pKB34 encoding the *yscW-lacZ* transcriptional fusion and the *yscWlcrF-lacZ* translational fusion were transformed into *Y. pseudotuberculosis* strains YPIII (wt) pAKH85 (pV, empty vector), YP356 (Δ*rnc*) pAKH85 (pV, empty vector) and the complementing strains YPIII (wt) pIVO20 (p*rnc*^+^) and YP356 pIVO20 (p*rnc*^+^). The transformants were grown at 37°C and alkaline phosphatase or beta-galactosidase activity was determined, respectively. Data represent the mean ± SD from three independent biological replicates. Significant differences were determined using Student’s t-test and indicated by asterisks (*P <0.05).

To investigate whether the absence of RNase III and PNPase affects the stability of the *lcrF* transcript, we determined the effects of RNase III on *lcrF* mRNA decay and measured the half-life of the transcript in the wildtype, the Δ*rnc,* Δ*pnp,* and Δ*rnc/*Δ*pnp* mutant by Northern blot analysis (Fig. **8A,B**). The half-life of the *lcrF* mRNA in the Δ*rnc* mutant was two-fold higher than in the wildtype. Loss of PNPase also increased *lcrF* stability although to a lesser extent, and the effect of both RNases on *lcrF* transcript stabilization seems to be additive (Fig. **8B**).

**Figure 8:**
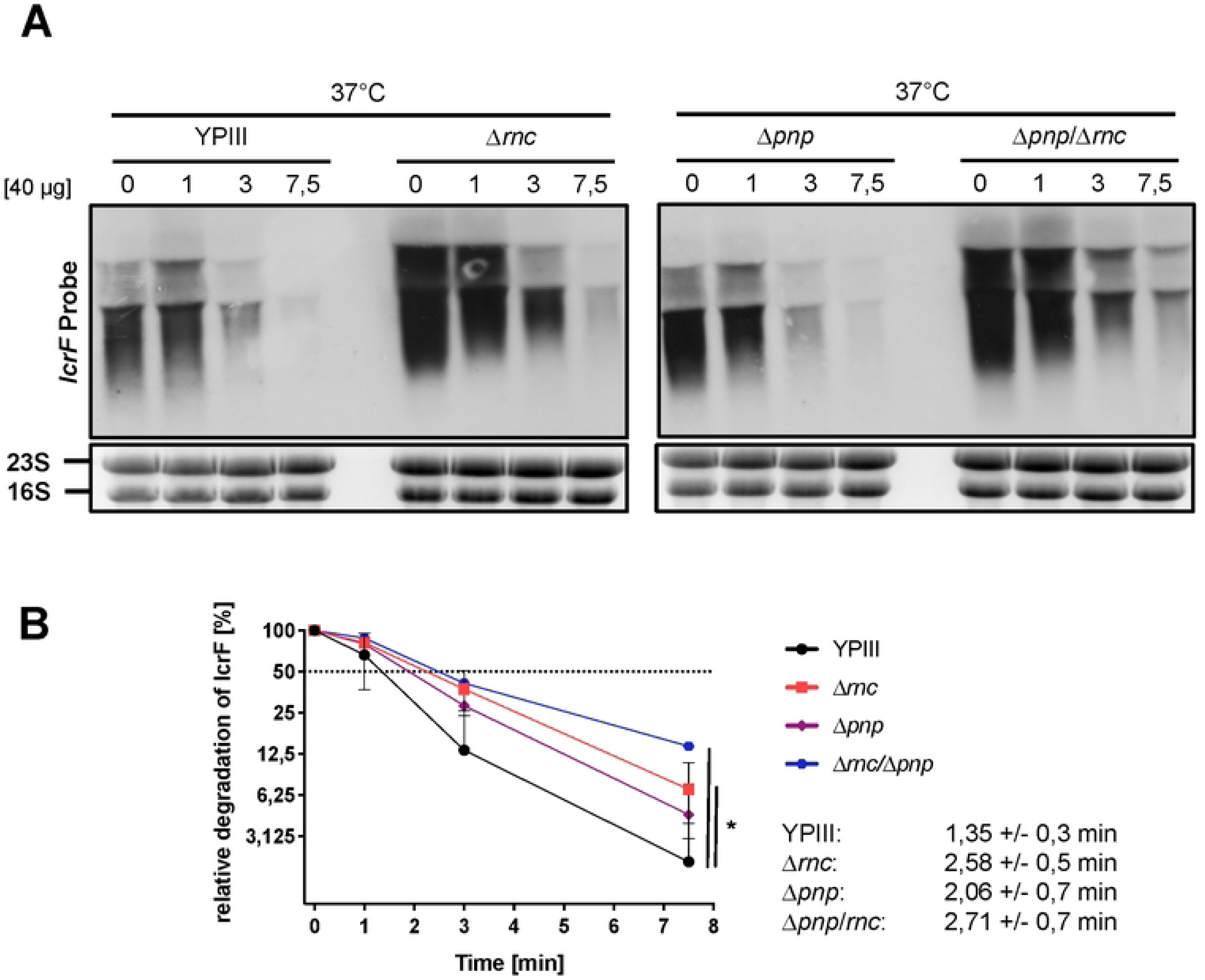
Influence of RNase III and PNPase on the stability of the *lcrF* transcript. *Y. pseudotuberculosis* strains YPIII (wt), YP356 (Δ*rnc*), YP139 (Δ*pnp*), and YP375 (Δ*rnc* Δ*pnp*) were grown at 37°C. Rifampicin was added to the cultures to stop transcription. Samples were taken before (0 min) and 1, 3 and 7.5 min after rifampicin addition. Total RNA of the samples was extracted and subjected to Northern blotting, using an *lcrF*-specific probe. 16S and 23S rRNA served as loading controls. A representative Northern blot for each strain is shown from three independent biological replicates. (**B**) The three independent biological replicates were used for the quantification of *lcrF* transcript levels with ImageJ software. 16S rRNA was used as a control for quantification. Relative transcript levels were plotted in Graphpad Prism 8 to determine the *lcrF* mRNA decay rate. Data represent the mean ± SD from three independent biological replicates. Significant differences were determined at 7.5 min using Student’s t-test and indicated by asterisks (*P <0.05).

Next, we investigated whether the RNases act directly on the *lcrF* mRNA, but according to our analysis, it is very unlikely that the *lcrF* transcript is a direct target of RNase III. Firstly, we did not identify potential RNase III cleavage sites (e.g. ∼22 nt double-stranded RNA segments including base-pairing preferences such as 5’ GAAAnnAAAG/CUUUnnUUUG 3’) within the *yscW-lcrF* transcript, which were identified in direct targets in previous surveys [49][50][53]. Only one longer double-stranded RNA segment without a potential RNase III cleavage sequence was identified in the 5’ untranslated region (5’-UTR) of the *yscW-lcrF* transcript. However, its presence in the 5’-UTR of the P*_yscW-lcrF_-lacZ* reporter fusion (pKB35) did not result in an expression change in the Δ*rnc* mutant compared to wildtype (Fig. **7B**). Secondly, no smaller-size degradation products of the *lcrF* transcript were detectable in the mRNA pool of the wildtype compared to the Δ*rnc* strain in gels with a high resolution (Supplementary Figure **S4**). Thirdly, although we detected a massive increase of reads mapping to the *lcrF* mRNA in our RNA-seq analysis when visualized with the same nucleotide score height (Fig. **6C**), an enlargement of the transcript profile resolution of the wildtype (from the nucleotide score height of 1000 to 100) revealed that the overall read pattern of *lcrF* was comparable with that of the Δ*rnc* mutant. No typical endoribonuclease-mediated processing events generating additional sharp/trimmed read ends in the wildtype were identified (Fig. **6D**) as described for direct mRNA targets [55,56]. These results indicated that RNase III is unlikely to target the *lcrF* mRNA directly at 37°C, but rather affects post-transcriptional regulators that have an overall influence on *lcrF* transcript levels, such as the effector protein YopD and the carbon storage regulatory system CsrABC [18].

### Mutual influence of RNase III, PNPase, and the translocon protein YopD

Besides its role as a structural component of the translocation pore of the T3SS for Yop delivery [57][58], YopD also acts as a negative post-transcriptional regulator that represses the synthesis of the master regulator LcrF and components of the Ysc-T3SS/Yop system under non-secretion conditions [18][59][60]. To test whether PNPase and RNase III affect YopD synthesis, we compared intracellular YopD levels between wildtype and RNase mutants.

The amount of YopD was strongly increased in the absence of RNase III at 37°C with a similar level observed in the wildtype at 37°C/-Ca^2+^ (Fig. **9A**). However, YopD upregulation does not explain the observed phenotypes of the Δ*rnc* mutant, as the presence of YopD was shown to reduce and not increase *lcrF* mRNA levels, and Ysc-T3SS/Yop synthesis [18][59][60]. On the contrary, YopD seems to be upregulated in the absence of RNase III as part of the entire Ysc-T3SS/Yop secretion complex due to higher LcrF levels (Fig. **5B**).

**Figure 9:**
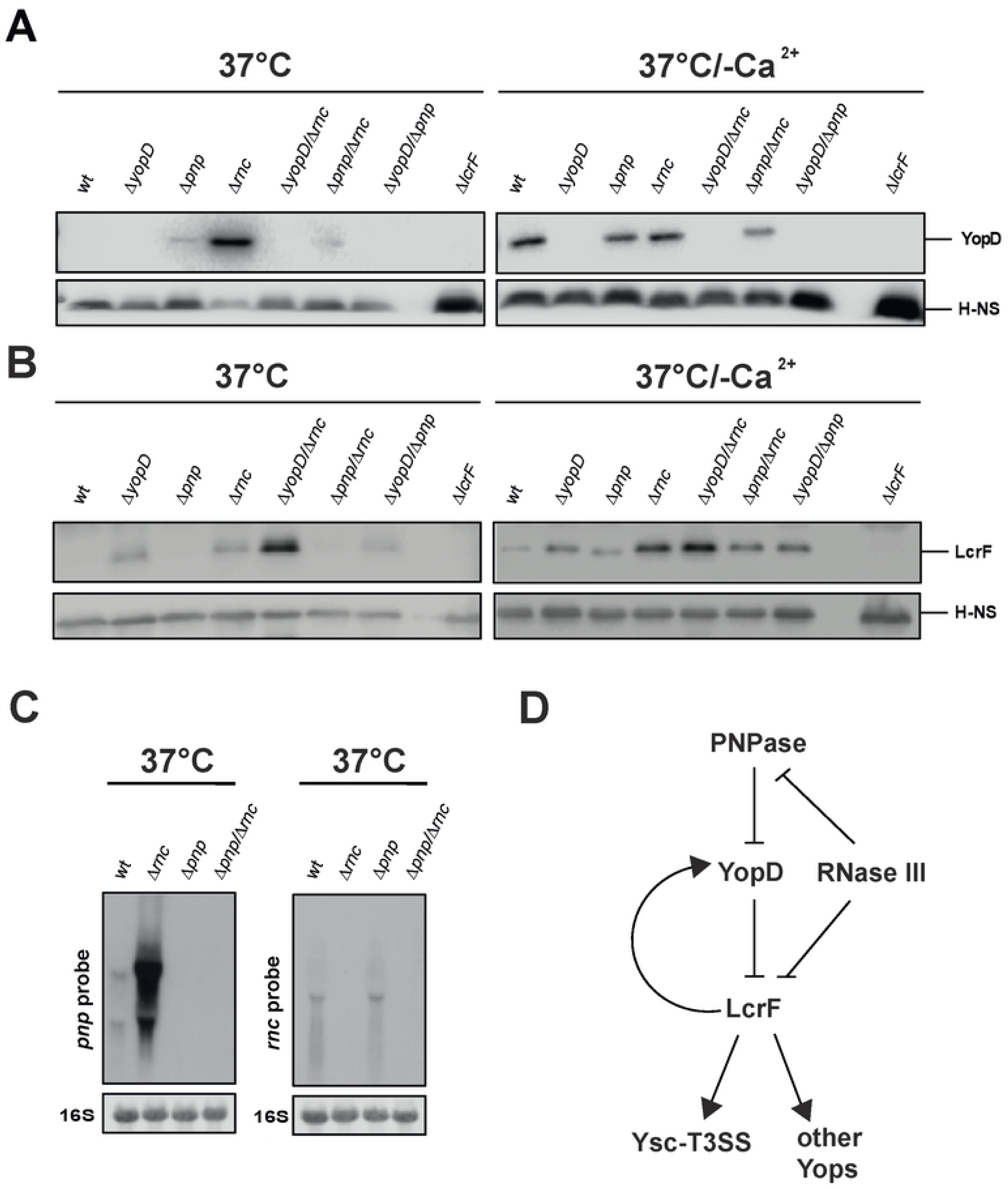
Influence of RNase III on the synthesis of YopD. *Y. pseudotuberculosis* strains YPIII (wt), YP91 (Δ*yopD*), YP179 (Δ*lcrF*), YP139 (Δ*pnp*), YP218 (Δ*yopD/*Δ*pnp*), YP372 (Δ*yopD/*Δ*rnc*), and YP375 (Δ*pnp/*Δ*rnc*) were grown at 37°C and 37°C/-Ca^2+^. Whole-cell extracts were prepared and separated by SDS-PAGE. Synthesized YopD (**A**) and LcrF (**B**) were detected by Western blotting using a YopD-detecting polyclonal antiserum. An antiserum against H-NS was used for loading control. (**C**) *Y. pseudotuberculosis* strains YPIII (wt), YP139 (Δ*pnp*), YP356 (Δ*rnc*) and YP375 (Δ*pnp/*Δ*rnc*) were grown at 37°C. The *pnp* (left) and the *rnc* (right) transcripts were detected by Northern blotting using specific probes. 16S RNA served as loading controls. (**D**) Scheme illustrating the control of LcrF and Ysc-T3SS-Yop synthesis by PNPase and RNase III.

Moreover, the amount of the transcriptional activator LcrF is similarly increased in the Δ*rnc* and Δ*yopD* mutant compared to wildtype, but the overall LcrF level is much higher in the Δ*rnc/*Δ*yopD* double compared to the single mutants (Fig. **9B**), indicating an additive effect. This shows that RNase III regulates *ysc*-T3SS/*yop* gene expression not via but rather together with YopD.

Loss of PNPase resulted also in the upregulation of YopD, but its influence was much less pronounced (Fig. **9A**). A comparison of the Δ*yopD* and the Δ*pnp/*Δ*yopD* mutant revealed no difference in LcrF levels, whereby the overall repressive effect of YopD was generally stronger than that of PNPase (Fig. **9B**). Together, this suggested that PNPase could act upstream of YopD.

Surprisingly, the production of LcrF and YopD did not further increase in the Δ*pnp*/Δ*rnc* double mutant as expected from the single mutation. In contrast, the Δ*rnc/*Δ*pnp* double mutant contained less YopD and LcrF than the Δ*rnc* single mutant (Fig. **9A,B**). This indicated an interplay of the RNases in which the loss of PNPase seems to counteract the upregulation of LcrF promoted by the loss of RNase III. It is known that RNase III cleaves within the 5’ end of the *pnp* transcript and initiates its rapid degradation in *E. coli*; as a result, PNPase mRNA levels are significantly increased in the *E. coli* Δ*rnc* mutant strain [61][62]. Significantly higher amounts of the *pnp* transcripts were also identified in the *rnc* mutant of *Y. pseudotuberculosis* grown at 37°C by the RNA-seq analysis (log_2_ = 2.2 (T1) 1.9 (T2); Data sheet **S4**) and by Northern blotting (Fig. **9C**), indicating a similar process in this pathogen. This suggests an interactive, regulatory pattern illustrated in Fig. **9D**, in which RNase III strongly represses LcrF production via a YopD-independent pathway, which is partially compensated by a second pathway controlling PNPase and YopD.

### Impact of RNase III on the Csr system

As the mechanism of how RNase III controls *lcrF* mRNA levels independently of YopD remained unclear, we tested whether RNase III influences the amount of CsrA. CsrA is another post-transcriptional regulator known to increase the stability of the *lcrF* transcript in response to host cell contact/secretion [18]. CsrA belongs to the carbon storage regulator (Csr) system together with the two regulatory RNAs CsrB and CsrC which sequester multiple CsrA dimers to eliminate their function (Fig. **10A**, right panel). Loss of RNase III did not influence the overall amount of CsrA (Fig. **10A**). However, expression of the counteracting CsrB and CsrC sRNAs was significantly reduced in the absence of RNase III (Fig. **10B**). This, in turn, increases the amount of unbound, active CsrA, which is known to strongly reduce the degradation of the *lcrF* mRNA [18]. The molecular mechanism of how CsrA stabilizes the *lcrF* mRNA is currently unknown [18]. However, as CsrA was found to enhance translation of the *lcrF* transcript [18], and a higher ribosome occupancy upon upregulation of translation was shown to protect mRNAs from RNase-mediated degradation [63][64], it seemed possible that increased availability of active CsrA in the Δ*rnc* mutant enhances translation and thus *lcrF* mRNA stability. Based on this assumption, we tested the influence of RNase III on the expression of a translational reporter fusion in which the *lacZ* gene is fused to codon 25 of the *lcrF* gene, harboring the intergenic RBS and the entire promoter region of the *yscW-lcrF* operon up to position -304 according to the transcriptional start site (Fig. **7C**). In contrast to the transcriptional fusions (Fig. **7A,B**), the translational reporter revealed a significant increase in the synthesis of the LcrF-LacZ fusion protein in the absence of RNase III (Fig. **7C**).

**Figure 10:**
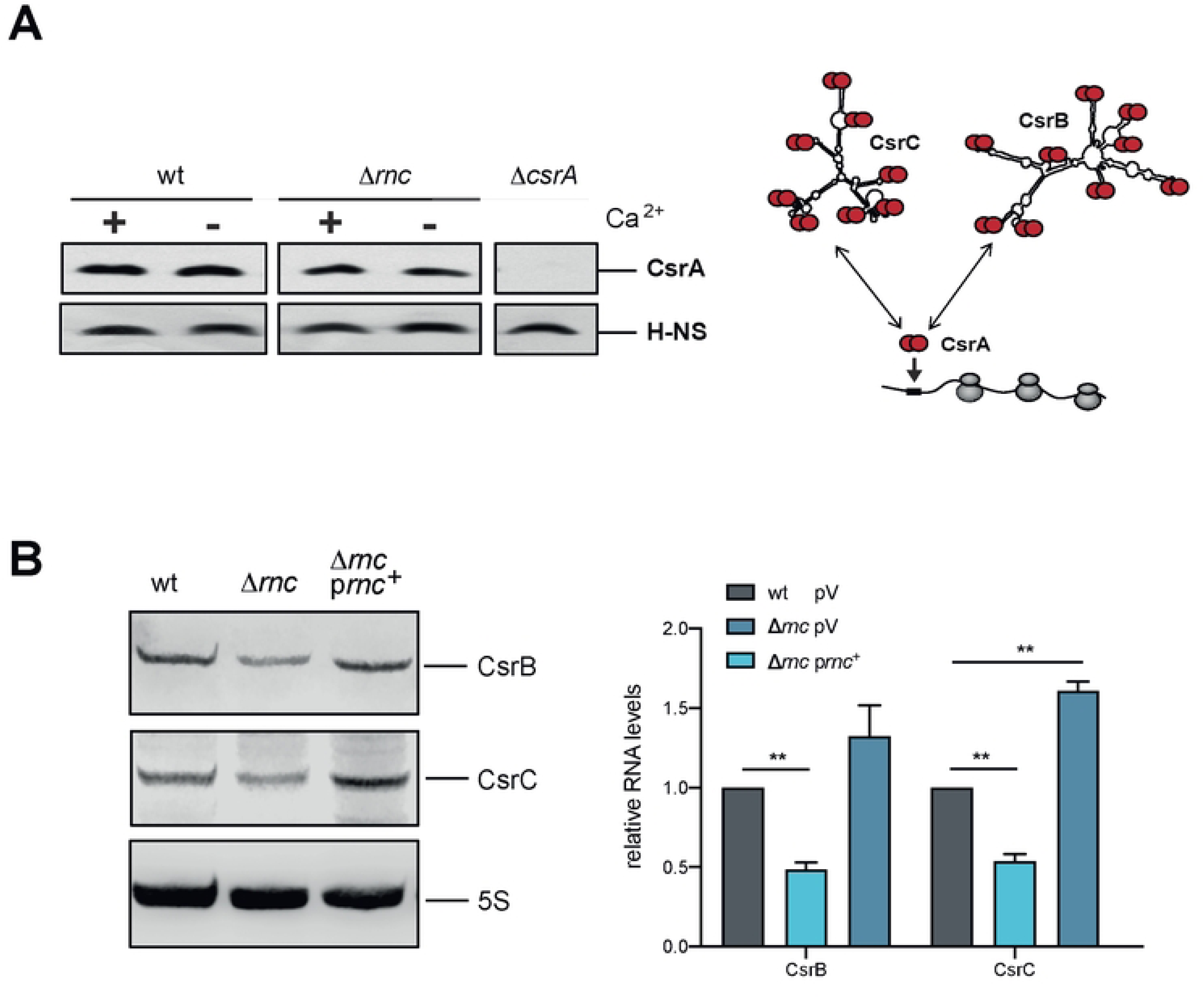
Influence RNase III on the carbon storage system components CsrA, CsrB and CsrC. (**A**) (Left) *Y. pseudotuberculosis* strains YPIII (wt) and YP356 (Δ*rnc*) were grown at 37°C in the presence (+) and absence of Ca^2+^ (-). Whole-cell extracts were prepared and separated by SDS-PAGE. Synthesized CsrA protein was detected by Western blotting using a polyclonal antiserum against CsrA. An antiserum against H-NS was used for loading control. (Right) Scheme of the interaction of the carbon storage regulatory systems components: the RNA binding protein CsrA and the CsrA-sequestering regulatory RNAs CsrB and CsrC. (**B**) (Left) *Y. pseudotuberculosis* strains YPIII (wt), YP356 (Δ*rnc*), and the complemented strains were grown at 37°C in the presence of Ca^2+^ (non-secretion conditions). Total RNA of the samples was prepared and CsrB and CsrC were detected by Northern blotting. (Right) Amounts of the CsrB and CsrC RNA were quantified by ImageJ and represent the mean ± SD from three independent biological replicates. Significant differences were determined using Student’s t-test and indicated by asterisks (**P<0.01).

### RNase III influence on the translational machinery

The increase of active CsrA could explain the Δ*rnc* phenotype as the CsrB and CsrC levels are reduced to approximately 50% in the RNase III-deficient strain (Fig. **10B**). However, additional identified expression changes of components of the translation machinery may have also an influence. Our RNA-Seq analysis comparing the transcription profiles of the wildtype and the Δ*rnc* mutant revealed that the transcript levels of 11 ribosomal proteins (r-proteins), as well as several translation initiation factors and rRNA/tRNA-modifying enzymes, were significantly altered with a log_2_ fold change of +/- ≤1 (Table **S1**). Among them are (i) the r-protein transcripts of *rpsJ* (S10, log_2_ = 3.059) and *rplC* (L3, 2.784) encoded by the S10 operon involved in rRNA and r-protein regulation [65]; (ii) the translational initiation factor IF-3 (InfC) (log_2_ of 1.02), an essential bona fide factor that promotes 30S initial complex formation, accurate tRNA selection, and fidelity of bacterial translation initiation [66][67], (iii) the 16S rRNA methyltransferase RsmG/GidB with the ability to enhance the loading of r-proteins and 30S ribosome assembly in the presence of the 5’-leader [68], and (iv) the 16S rRNA-processing protein RimM assisting in the assembly of the 30S subunit, and promoting translation efficiency [69][70]. The higher abundance of these translation factors in the Δ*rnc* mutant could also increase (or assist CsrA-mediated activation of) *lcrF* translation and thus *lcrF* mRNA stability.

### RNase III-mediated global reprogramming of the *Y. pseudotuberculosis* transcriptome in response to virulence-relevant conditions

Based on our results, we assumed that loss of RNase III has a global influence on the gene expression pattern of the *Yersinia* chromosome and the virulence plasmid. To obtain more information about the overall influence of RNase III, we performed transcriptomic profiling by strand-specific RNA sequencing followed by a differential expression analysis (DESeq) by comparing sequence reads from strand-specific libraries of *Y. pseudotuberculosis* wildtype and the Δ*rnc* mutant strain. For this purpose, the bacteria were grown at 25°C (T0) (environmental, non-T3SS expression conditions), and then shifted to 37°C (host, T3SS non-secretion), and 37°C/-Ca^2+^ (host, T3SS secretion conditions) for 1 h (T1) and 4 h (T2). 1 h was chosen as this is the earliest time point at which an LcrF/Yop upregulation can be detected, and 4 h was chosen for maximal T3SS induction indicated by the strong expression of the activator LcrF, and synthesis of the Yops (Fig. **S5**). From each library, between 2-3 million cDNA reads were generated and mapped to the YPIII genome sequence (NC_010465) and the pYV virulence plasmid (NC_006153) (Dataset **S1**). The global gene expression profiles of the different samples were distinct, and profiles of the three biological triplicates clustered together (Fig. **S6**).

The detailed analysis of the transcriptome revealed that approximately >99% of the reads of the *Y. pseudotuberculosis* YPIII (wt) and the Δ*rnc* mutant grown at 25°C mapped to the chromosome (4,7 Mb, 4,250 genes). The read distribution was only slightly changed in the wildtype when the bacteria were shifted to 37°C. Under these conditions, more reads were found to map to the virulence plasmid pYV (99 genes, 66 kb) at T1 and T2. However, this still accounts for less than 3% of the total reads (Fig. **11A**) (Datasets **S2**). This changed significantly under secretion conditions (induced by the depletion of Ca^2+^). Approximately 56% (T2) of the reads were assigned to the pYV, and only 44% (T2) were mapped to the chromosome. This indicated a major shift of the transcriptome from chromosomally- to virulence plasmid-encoded genes. Most strikingly, a similar shift was observed in the Δ*rnc* mutant grown at 37°C under non-secretion conditions (+Ca^2+^) (Fig. **11A**, Datasets **S3**). About 47% (T2) of the entire transcripts originated from the virulence plasmid, which resembled the situation in the wildtype under secretion conditions (-Ca^2+^). This is particularly striking because the virulence plasmid represents only 1.5% of the entire *Y. pseudotuberculosis* genome.

**Figure 11:**
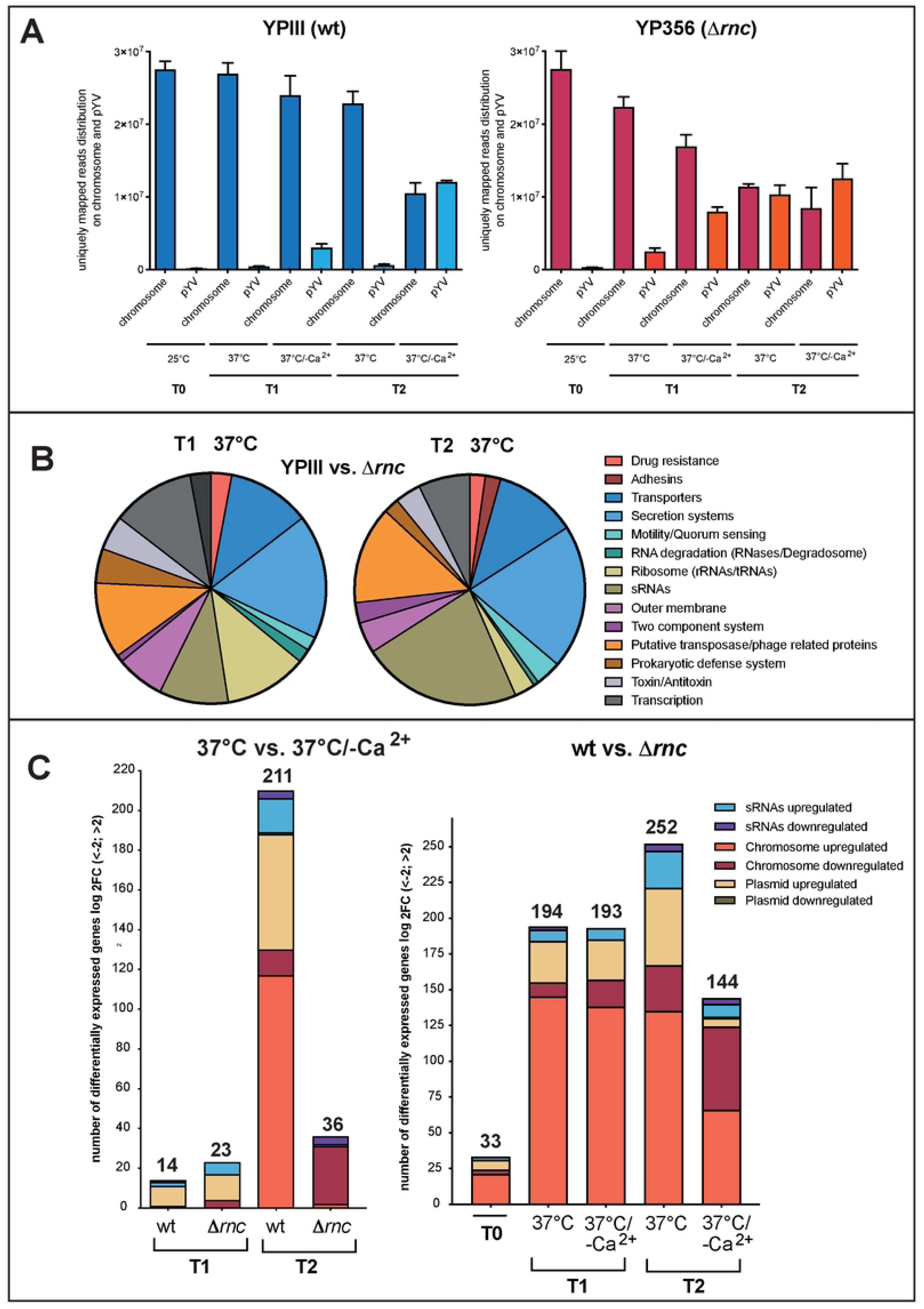
Global influence of RNase III on the *Y. pseudotuberculosis* transcriptome. *Y. pseudotuberculosis* strains YPIII (wt), and YP356 (Δ*rnc*) were grown at 25°C, and at 37°C in the presence and absence of Ca^2+^ for 1 h (T1) or 4 h (T2). Total RNA of the different strains was prepared and transcriptional profiling by strand-specific RNA sequencing followed by a differential expression analysis (DESeq) by comparing sequencing reads of wildtype and the Δ*rnc* mutant strain was performed. (**A**) RNA-Seq read the distribution on the chromosome and the virulence plasmid (pYV). RNA-Seq reads which uniquely mapped to the chromosome or pYV under the different tested conditions are shown. (**B**) Influence of RNase III on global pathways at T1 and T2 at 37°C. Differentially expressed genes between wildtype and the Δ*rnc* mutant with a log2FC cut-off of -2/+2 and P-value ≤ 0.05 were used for the analysis of global biological pathways based on the categories established according to the KEGG database. (**C**) Quantitative analysis of differentially expressed genes between wildtype and the Δ*rnc* mutant. Bar plots show the absolute number of differentially expressed genes (left panel) after Ca^2+^ depletion after 1 h (T1) or 4 h (T2) of the wildtype (wt) or the Δ*rnc* mutant, or (right panel) between the wildtype (wt) and the Δ*rnc* mutant at 25°C (T0), before the shift to 37°C for 1 h (T1) or 4 h (T2) in the presence or absence of Ca^2+^.

To identify which global pathways are particularly affected by the absence of RNase III at 37°C under non-secretion conditions, we summarized the genes up- or downregulated (log2-fold change (log2FC) ≥ +/- 2, p-value ≤ 0.05) in groups based on their association with the pathway categories found in the KEGG Orthology database. (Fig. **11B**, Dataset **S4**). We found that mostly pYV-encoded genes and several regulatory RNAs are upregulated in the wildtype and the Δ*rnc* mutant 1 h (T1) after the shift from non-secretion (37°C) to secretion conditions (37°C/-Ca^2+^) (Fig. **11C**, left panel, Datasets **S2**, **S3**). This pattern changed after 4 h (T2) when an additional upregulation of chromosomally encoded transcripts mostly for metabolic enzymes and transporters was observed in the wildtype (likely to compensate for the energetic burden caused by T3SS/Yop synthesis), but not in the Δ*rnc* strain (Fig. **11C**, left panel, Datasets **S2**, **S3**).

Our results further showed that loss of RNase III at 25°C (T0: 33 genes) was rather moderate, whereas a shift to 37°C had a significant impact on the transcripts of numerous genes compared to wildtype (37°C, T1: 194 genes, T2: 252 genes) (Dataset **S4**, Fig. **11C**, right panel). This indicated that the influence of RNase III is strongly temperature-dependent. At 37°C, many more metabolic gene transcripts were detected in the wildtype (Dataset **S4**, Fig. **11C**, right panel). This may explain the generally lower fitness of the RNase III-deficient strain at this growth condition (Fig. **1A**).

Notably, the abundance of almost all pYV-encoded Ysc-T3SS/Yop-encoding transcripts, including the transcripts of the *yscW-lcrF* operon was strongly increased in the Δ*rnc* mutant at 37°C compared to wildtype at 37°C (Table **S2**, Dataset **S2,** Fig. **11B**, right panel), but it was very similar to the wildtype at 37°C/-Ca^2+^ (Table **S2**, Dataset **S2**, Fig. **11C**, right panel). Based on this analysis, we hypothesized that secretion-relevant genes are already induced under non-secretion conditions at 37°C in the absence of RNase III. To prove this assumption, we compared transcriptome changes of the wildtype upon the shift from non-secretion to secretion conditions (YPIII, 37°C/+Ca^2+^ vs. 37°C/-Ca^2+^) (Dataset **S2**) with changes of the wildtype and the Δ*rnc* mutant at 37°C under non-secretion conditions (YPIII, 37°C/+Ca^2+^ vs. Δ*rnc*, 37°C/+Ca^2+^) (Dataset **S4**). Notably, of the 28 pYV-encoded transcripts which were found in a higher abundance in the Δ*rnc* mutant at 37°C at T2, 27 were also increased under secretion conditions in the wildtype to almost the same magnitude. This indicated that *lcrF* and T3SS/*yop* expression upon host cell contact could be triggered by a reduction of functional RNase III during the infection.

We further found that besides pYV-encoded factors, only a few other chromosomally-encoded transcripts were induced in the Δ*rnc* mutant at 37°C at T1; among them are several translation-relevant factors, e.g. the ribosome-association factor Rai/YfiA (log2FC=1.5), the ribosomal protein S2/RpsB (log2FC=1.3), and the translation initiation factor IF3 (log2FC=1). These components of the translational machinery could also promote/assist in the global reprogramming of the *Yersinia* transcriptome in the absence of RNase III.

## Discussion

Gram-negative bacteria possess roughly 20 RNases which control RNA turnover and maturation, and this plays an important role in many virulence-relevant processes [4][9][10]. RNases can possess specificity towards certain RNA types (e.g. sRNA or dsRNA) and their activity can also depend on the intrinsic folding of the RNA, length of the transcript, bound ribosomes, and regulatory proteins [4][9][10]. Among them are the highly conserved 3’->5’ exonuclease PNPase and the endoribonuclease RNase III. In this study, we demonstrate that both RNases are global regulators in *Y. pseudo-tuberculosis* with multiple cellular targets, including virulence factors, such as the type III secretion systems affecting motility and Yop secretion.

Loss of PNPase had a moderate influence on the expression of T3SS/*yop* genes, and did not severely affect flagella T3SS-mediated motility. In contrast, the absence of RNase III abolished motility and flagella synthesis (similar to *E. coli rnc* mutants, which have no more than 1 in 1000-10,000 cells with sufficient flagella [71]) and led to a strong increase of T3SS injectisome formation and Yop protein synthesis and secretion under normally non-inducing conditions (37°C/+Ca^2+^). The *ysc*-T3SS/*yop* expression pattern of the *rnc* strain was very similar to that of the wildtype under secretion conditions (37°C/-Ca^2+^), suggesting that loss of RNase III triggered a process equivalent to host-cell contact/Ca^2+^ depletion.

A detailed analysis assigned to identify the underlying molecular mechanism revealed that Ysc-T3SS/Yop induction in the Δ*pnp* and Δ*rnc* mutant occurred through the upregulation of the AraC-type master activator LcrF of the *ysc*-T3SS/*yop* genes. As expression of the *lcrF* gene is part of a complex regulatory network, implicating multiple transcriptional regulators (YmoA, H-NS, IscR, RcsB), post-transcriptional regulatory elements (e.g. a fourU RNA thermometer, RNA binding proteins YopD and CsrA) [18][31][32] as well as plasmid-copy number control systems (RepA, CopA/B) [47], we tested the influence of both RNases on different control levels of LcrF synthesis.

We found that the loss of both RNases did not affect the transcription of the *lcrF* gene and the copy number of the virulence plasmid, but led to increased *lcrF* mRNA levels/stability. However, against our first assumption, the *lcrF* mRNA is most likely not a direct target of the RNases. RNase III is known to target intramolecular and intermolecular double-stranded RNA segments located mostly 5’ or 3’ to a coding sequence [49][50]. However, no sequences with similarity to the described base-pairing preferences of RNase III cleavage sites could be identified within the *yscW-lcrF* transcript. Moreover, no characteristic RNase-mediated degradation products and no typical sequencing read pattern changes of the *lcrF* mRNA due to processing by RNase III or PNPase were detectable, as described for targeted mRNAs of other bacteria [55,56].

This suggested that the observed RNase-mediated influence on *lcrF* transcript levels may be an indirect consequence of the control of regulators known to affect the stability of the *lcrF* mRNA. One such regulator is YopD, whose intracellular presence in the absence of host cell contact/secretion was shown to destabilize the *lcrF* transcript [18]. The comparative analysis of Δ*pnp,* Δ*rnc,* and Δ*yopD* single and double mutants and their influence on LcrF synthesis revealed a complex control circuit in which RNase III, PNPase, and YopD are tightly interconnected and mutually affect their synthesis (Fig. **12**). In this network, PNPase reduces the stability of the *lcrF* mRNA, most likely through downregulation of YopD. The overall influence of PNPase on *lcrF* transcript levels via YopD is rather moderate. This can be explained by feedback control (Fig. **12**), as the loss of YopD also resulted in a significantly lower abundance of *pnp* mRNAs [18]. Similar to PNPase, RNase III reduces the stability of the *lcrF* transcript, but its overall influence is much more pronounced. Loss of the *rnc* gene led to much higher *lcrF* mRNA levels, and this effect could be further enhanced by the deletion of the *yopD* gene. We also observed that *pnp* transcript levels were significantly increased in the Δ*rnc* mutant, indicating that RNase III cleaves within the *pnp* transcript and initiates its degradation as described for *E. coli* [61][62]. These observations suggested two separate pathways in which (1) RNase III has a very strong and dominant negative influence on the *lcrF* mRNA independent of YopD, and (2) RNase III exerts a YopD-dependent compensatory effect on *lcrF* mRNA stability via the repression of the PNPase (Fig. **12**). The second pathway may have evolved to balance the negative influence of RNase III on *lcrF* mRNA stability in the absence of host cell contact, whereby an additional upregulation of LcrF production is still possible upon YopD secretion.

**Figure 12:**
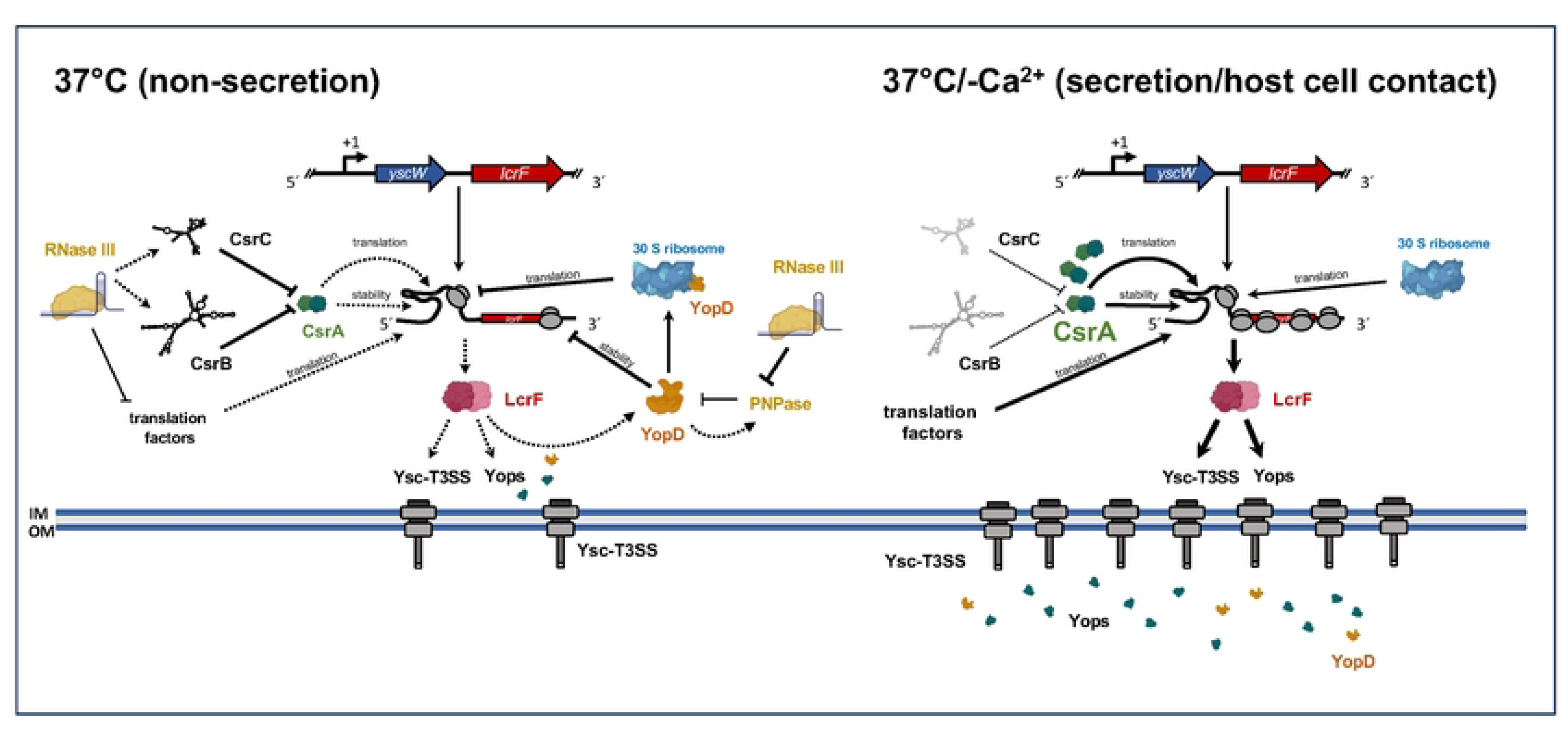
Scheme of PNPase and RNase III-dependent synthesis of LcrF and the T3SS/Yops under non-secretion and secretion conditions. Regulatory network of the LcrF-T3SS/Yop system is illustrated under non-secretion (left panel) and secretion condition (right panel). Strong control is indicated by a thick solid arrow for activation or T for inhibition, low level of control is indicated by dashed arrows for activation or T for inhibition.

The exact molecular mechanism of how RNase III promotes the destabilization of the *lcrF* mRNA by pathway (1) is still unknown. However, the results of this study indicate that this may (at least in part) occur through the control of the RNA-binding protein CsrA (Fig. **12**), a post-transcriptional regulator of the Csr system, known to stabilize the *lcrF* mRNA and activate its translation by binding to the *lcrF* 5’-UTR [18]. Here, we show that loss of RNase III has no effect on the overall levels of CsrA but leads to a significant decrease in the regulatory RNAs CsrB and CsrC. As both Csr RNAs were shown to bind and sequester multiple CsrA dimers to block their activity [72], a higher amount of unbound, active CsrA is expected in the Δ*rnc* mutant. An increase in active CsrA could explain (i) higher stability of the *lcrF* mRNA, and (ii) higher expression of the translational *lcrF-lacZ* fusion in the absence of RNase III. It is known that CsrA interacts directly with the 5’-untranslated region of the *lcrF* transcript and enhances *lcrF* mRNA translation [18] which could also increase *lcrF* mRNA stability. In fact, studies with *E. coli* showed that translation initiation regions (5’-UTR + first 11 codons of the downstream gene) that dictate high translation initiation rates (e.g. high RBS indexes and/or absence of strong local secondary structures) recruit more ribosomes to the mRNA, and the high ribosome occupancy can protect the transcript from degradation by RNases [73][74].

It is also possible that other translation-relevant factors that were shown to be differrentially expressed in the Δ*rnc* mutant support this CsrA function or enhance translation initiation separately, which could increase ribosome coverage and thus *lcrF* transcript stability. Among these factors are 11 ribosomal proteins, and the essential bona fida translation initiation factor IF-3 (InfC) promoting 30S initial complex formation [66][67]. Moreover, transcripts of the 16S rRNA methyltransferase RsmG/GidB and the 16S rRNA-processing protein RimM were significantly increased in the absence of RNase III which both also have a positive influence on translation initiation, e.g. by subverting problems that occur during the *in vivo* assembly of the 30 S ribosome [68][69][70][75] or by guiding the folding of the rRNAs to facilitate the formation of productive r-protein-RNA interactions [76]. In contrast, the transcript of the ribosome-associated inhibitor A (RaiA/YfiA), which accumulates in the stationary phase, and stabilizes 70S ribosomes in an inactive state [77], was less abundant in the absence of RNase III.

The observed changes regarding the translation-relevant factors could be a consequence of defects in rRNA processing associated with the loss of RNase III. It is well known that RNase III initiates maturation of the ribosomal RNAs encoded on the pre-rRNA transcript by an initial series of processing events, which are subsequently cleaved and trimmed by other ribonucleases [78–81][82]. In RNase III-deficient bacteria 30 S rRNA precursors (including 16S, 23S, 5S rRNA, and tRNA sequences) are produced [78–81][83]. The 30 S rRNA can be processed by alternative maturation pathways. However, this involves the formation and incorporation of not fully matured rRNAs (prRNAs) into the ribosomes, which alter their overall activity [84][82,85,86]. The RNase III-deficient bacteria may indirectly compensate for these alterations by the upregulation of translation factors. However, the influence of RNase III on translation factors could also be direct. Multiple RNase III *in vivo* cleavage sites were recently found in mRNAs encoding ribosomal proteins, translation elongation factors, and enzymes involved in RNA modification in *E. coli* [52][87]. This indicated that the role of RNase III in the control of translation is much larger than expected, and could support global reprogramming of bacterial gene expression, e.g. upon host cell contact, by altering the translational machinery.

Apart from the *ysc*-T3SS/*yop*-related alterations, the RNase III-deficient strain is characterized by a wide range of transcriptional changes, including mRNAs and sRNAs across all categories (e.g. metabolism, additional virulence traits, responses to environmental stresses, genetic/environmental information processing). Among them are several other RNA-controlling factors, i.e. small regulatory RNAs encoded on the virulence plasmid (e.g. Ysr291-293 overlapping with the *lcrV-lcrH-yopD* transcript), which could also affect *lcrF* mRNA stability.

Approximately 10% of all mRNAs are affected by the introduction of the Δ*rnc* mutant in *Yersinia*. This is similar to other bacteria, whereby the impact on individual genes varies substantially among the species [88][89][90]. Our comparison further revealed that only 1-2% of the total uniquely mapped reads were generated from the virulence plasmid pYV at conditions reflecting the environment or initial stages of the infection. However, this drastically changed under Ysc-T3SS/Yop-inducing conditions mimicking host cell contact. Although only approximately 1% of the genetic information of *Y. pseudotuberculosis* is encoded on pYV, it accounts for more than 50% of the transcripts detected under secretion conditions, whereas the ratio of the chromosomally-encoded transcripts was drastically reduced (from >99% to less than 50%). This may also explain the growth arrest associated with the induction of Ysc-T3S-mediated secretion as the abundance of many transcripts important for metabolism and physiological traits is affected. It also became apparent that in particular mRNAs of pYV were already upregulated in the Δ*rnc* mutant under non-secretion conditions. This suggested that reprogramming observed under secretion conditions in the wildtype might be triggered by a reduction of functional RNase III. However, no change in *rnc* transcript levels was detectable upon a shift from non-secretion to secretion conditions. In this respect, RNase III is either not the superior trigger of LcrF/Ysc-T3SS/Yops or it is modulated on the post-transational level.

In summary, several aspects of this study indicate that RNase III-mediated control of LcrF and Ysc-T3SS/Yop production is (i) linked to components or regulators, such as CsrA and YopD, cooperating with the translational machinery, and (ii) accompanied by a global reprogramming of gene expression. Interestingly, translation has also previously been reported to play a role in LcrF, and Ysc-T3SS/Yop synthesis in another context. For instance, it was demonstrated that the RNA-binding translocator protein YopD associates with the 30S ribosomal particles in an LcrH-dependent manner and that this prevents translation initiation of its target mRNAs, including *lcrF* and the *yop* transcripts [91]. Hence, RNase III-mediated alterations of the composition and activity of the ribosome could in turn also modulate the interaction of YopD with the ribosome and affect its function as a regulator of *lcrF* mRNA translation and stability.

Our current understanding of the role of RNases in the complex post-transcriptional control of *Yersinia* virulence is still in its infancy. However, it is clear, that their involvement not only enables *Yersinia* to rapidly adjust its gene expression profile to efficiently block immune cell attacks, but it also allows the bacteria to overcome accompanying energetic and stress burdens and to rapidly resume its genetic pre-attack program after a successful defense.

## Experimental Procedures

### Cell culture, media, and growth conditions

Overnight cultures of *E. coli* were routinely grown in LB at 37°C, *Yersinia* strains were grown at 25°C or 37°C in LB (Luria Bertani) broth. For the analysis of Yop and LcrF synthesis and secretion under secretion and non-secretion conditions, overnight cultures of the *Yersinia* strains grown at 25°C were diluted in LB to an OD_600_ of 0.2, grown for 2 h at 25°C and then either further grown at 25°C, or shifted to 37°C in the presence or absence of the Ca^2+^ chelators 20 mM NaOx and 20 mM MgCl_2_ (37°C/-Ca^2+^). If necessary, antibiotics were added at the following concentrations: carbenicillin 100 µg ml^-1^, chloramphenicol 30 µg ml^-1^, kanamycin 50 µg ml^-1^, rifampicin 1 mg/ml and Triclosan 20 µg ml^-1^.

### Strain and plasmid constructions

All DNA manipulations, restriction digestions, ligations, and transformations were performed using standard genetic and molecular techniques [92][93]. Plasmid DNA was isolated using QIAprep Spin Miniprep Kit (Qiagen) or Nucleospin Plasmid Kit (Macherey-Nagel). DNA-modifying enzymes and restriction enzymes were purchased from New England Biolabs. The oligonucleotides used for amplification by PCR, sequencing, and primer extension were purchased from Metabion and Eurofins. PCRs were done in a 50 µl mix for 30 cycles using Phusion High-Fidelity DNA polymerase (New England Biolabs). Purification of PCR products was routinely performed using the QIAquick PCR Purification Kit (Qiagen) or the NucleoSpin Gel and PCR Clean-up Kit (Macherey-Nagel). All constructed plasmids were sequenced by Microsynth AG (Balgach, Switzerland).

Strains and plasmids used in this study are listed in Supplement Table **S3** and primers for plasmid generation are listed in Supplement Table **S4**. Plasmid pIVO13 was constructed by the insertion of a PCR fragment harboring the regulatory region of the encoding region of *yopE*. The fragment was amplified from total DNA of *Y. pseudotuberculosis* strain YPIII using primer set VIII193/VIII194 cloned into the *Bam*HI/*Xho*I sites of plasmid pPCH1, replacing the full-length coding sequence of *yopE*. The resulting plasmid pIVO13 was sequenced using primers I984 and V824. Plasmids encoding the *rnc* and *pnp* genes for complementation analyses were constructed by amplification of DNA segments including the respective native promoter identified by Nuss *et al*. [36], the coding sequence and the 3’-UTR of the RNase gene using primer pairs VIII365/VIII387 for *rnc* and VIII636/VIII/637 for *pnp* (Table **S4**). The generated PCR fragments were cloned into plasmid pAKH85 generating pIVO20 and pIVO21, and their sequence was confirmed using primer VIII385 and VIII388. Plasmid pTT15 was constructed by ligation of the *tet* promoter and the MCS with primers VI527 and II143 into the *Bam*HI and *Not*I sites of pFU86. The *yscW-lcrF* promoter region was amplified with primers IX729 and IX730 and ligated into the *Bam*HI and *Kpn*I sites of pTT15, generating pMV53.

### Construction of the Y. pseudotuberculosis deletion mutants

The *Y. pseudotuberculosis* strains used in this study are derived from wild-type strain YPIII. All deletion mutants were generated via homologous recombination. The mutant strains YP356, YP372, and YP375 in which the RNase III gene *rnc* was deleted were constructed in a three-step PCR. First, the *Yersinia* genomic DNA was used as a template to amplify 500-bp regions flanking the *rnc* gene. The upstream fragment was amplified with primer pairs (VII882/VII881) of which the reverse primer contained an additional 20 nt at the 5’-end which were homologous to the upstream region of the *rnc* gene. The downstream fragment was amplified with primer pairs (VII879/VII880) of which the forward primer contained additional 20 nt at the 5’-end which were homologous to the downstream region of *rnc*. Subsequently, a PCR reaction was performed with the forward primer of the upstream fragment and the reverse primer of the downstream fragment using the upstream and downstream PCR products as templates. The PCR fragment was digested with *Sac*I and was ligated into the *Sac*I site of the suicide plasmid pAKH3 generating plasmid pIVO11. The resulting plasmid was sequenced using primers III981/III982. Next, the plasmid was transformed into S17-1λpir and was transferred into *Y. pseudotuberculosis* YPIII, YP91 (Δ*yopD*), and YP139 (Δ*pnp*) via conjugation. Chromosomal integration of the plasmid was selected by plating on LB supplemented with kanamycin. Single colonies were tested for the correct integration of the plasmid via colony PCR using primer pair VII366/VII367. Mutants were subsequently grown on LB agar plates containing 10% sucrose. Single colonies were tested for growth on carbenicillin and sequenced using primer pair VII366/VII367.

### RNA isolation and Northern blotting

Bacterial cultures were grown under the desired conditions. The bacteria were pelleted by centrifugation for 1 min at 16000 x g. The pellets were resuspended in 0.2 volume parts of stop solution (5% water-saturated phenol, 95% ethanol) and immediately snap-frozen in liquid nitrogen. After thawing on ice, bacteria were pelleted for 1 min at 16,000 x g at 4°C. The pellet was resuspended in lysozyme solution (50 mg lysozyme/ml TE-buffer) and incubated for at least 5 min at RT. The total RNA of lysed bacteria was isolated with the ‘SV Total RNA Isolation System’ (Promega, USA) according to the manufacturer’s instructions. RNA concentration and quality were determined by measurement of A_260_ and A_280_.

For the analysis of *lcrF* transcripts total cellular RNA (5-30 μg) was separated on MOPS agarose gels (1.2%) or 8% urea acrylamide (RNA <250 nt), and transferred in 10 x SSC buffer (1.5 M NaCl, 0.15 M sodium citrate pH 7.0) for 1.5 h at a pressure of 5 cm Hg to a positively charged nylon membrane by vacuum blotting or by semi-dry blotting (250 mA, 2.5 h) and UV cross-linked. Prehybridization, hybridization to DIG-labeled *lcrF* PCR probes, and membrane washing were conducted using the DIG luminescent Detection kit (Roche) according to the manufacturer’s instructions. The *lcrF* transcripts of interest were detected with a DIG-labeled PCR fragment (DIG-PCR nucleotide mix, Roche) amplified with primer pairs for the individually detected according to the manufacturer’s instructions. (Table **S4**). For the detection of the CsrB and CsrC RNAs and the 5S and 16S control rRNA, fluorophore-coupled DNA oligonucleotide primers purchased from Integrated DNA technologies were used (Table **S4**). The primers were incubated with the membrane in a final concentration of 15 nM for CsrB and CsrC and 5 nM for 5 S rRNA in church moderate buffer (10 mg/ml BSA, 500 nM Na_2_HPO_4_, 0.34% (w/v) H_3_PO_4_, 15% (v/v) formamide, 1 mM EDTA, 7% SDS, adjusted to pH 7.2) for 6 h. The membrane was washed with 2 x SSC and 0,2% SDS for 5 min and the RNAs were detected at 700 nm (CsrB, CsrC) or 800 nm (5 S rRNA) with the Odyssey Fc Imaging system (Li-COR).

### RNA isolation for RNA-seq

To yield intact high-quality RNA for RNA-seq analyses, hot phenol RNA extraction was performed. The bacteria were harvested by centrifugation at 4,000 rpm and the bacterial pellets were snap-frozen in liquid nitrogen. Bacterial pellets were resuspended in 1/8 volume of RNA resuspension buffer (0.3 M saccharose, 0.01 M NaOAc) on ice. An equal volume of RNA lysis buffer (2% SDS, 0.01 M NaOAc pH 4.5) was added, mixed, and incubated at 65°C for 90 sec. An equal volume of pre-heated (65°C) saturated aqua-phenol was added, mixed with the lysate, incubated for 3 min at 65°C, and then immediately frozen in liquid nitrogen. Subsequently, the samples were centrifuged at 16,000 x g for 10 min at room temperature and the aqueous layer was extracted and the procedure was repeated three times. For the final extraction of phenol, an equal volume of chloroform/isoamyl alcohol (24:1) was added, mixed, and centrifuged for 10 min at 16,000 x g at room temperature. The RNA in the aqueous layer was precipitated by adding 1/10 volume of 3 M sodium acetate (pH 4.5) and 2.5 volumes of ice-cold ethanol at -20°C. The RNA was recovered by centrifugation at 16,000 x g for 30 min at 4°C, the pellets were washed at 4°C with ice-cold 70% (v/v) ethanol two times and the RNA pellets were air-dried and resuspended in nuclease-free water.

### RNA stability assay

**(A)** *Y. pseudotuberculosis* strains YPIII, YP139 (Δ*pnp*), YP356 (Δ*rnc*), and YP375 (Δ*pnp,* Δ*rnc*) were grown at 37°C. Subsequently, rifampicin was added in a final concentration of 1 mg/ml to block transcription, and samples were taken in intervals before (0), and 1, 3, and 7.5 min after the addition of rifampicin. To finally analyze the decay rate of the RNA transcripts, Northern blots were performed with an *lcrF*-specific probe as described [18], and the detected relative amount of RNA in each sample was determined using the Odyssey Fc Imaging system and software (LiCor).

### Quantitative PCR (qPCR) analysis of the copy number of pYV

The copy number of the *Y. pseudotuberculosis* virulence plasmid pYV was determined using quantitative PCR (qPCR). Total DNA of YPIII or its derivatives, grown at the indicated conditions, was isolated by the addition of chloroform:isoamylalcohol (24:1). The sample was mixed, centrifuged at 16,000 x g and the supernatant was mixed with four volumes of 100% ethanol. The precipitated DNA was pelleted by centrifugation (16,000 x g) and the pellet was resuspended in nuclease-free water. For the qPCR, three primer pairs for chromosomal genes (*glnA* (VII922/VII923), *rpoB* (VII924/VII925), YPK_3178 (VII926/VII927) and two primer pairs for plasmid genes (*yscM* (VII918/VII919), *repA* (VII920/VII921)) were chosen adapted from Wang *et al*. [47]. 1 ng/μl of total DNA was amplified using the primer sets in SensiFAST SYBR No-ROX (Bioline). qPCR was realized in technical duplicates in the Rotor-Gene Q real-time PCR cycler (Qiagen). Two reactions with nuclease-free water without DNA were used as controls. A melting curve was employed to ensure specificity. The plasmid copy number of the strains was calculated as described by Wang *et al*. [47].

### RNA sequencing (RNA-seq)

**(A)** *Y. pseudotuberculosis* YPIII and the isogenic *rnc* mutant YP356 were grown in LB medium to exponential phase (OD_600_ 0.5). The cultures were split and grown at 25°C (T0), 37°C, or 37°C in the presence of 20 mM Na_2_C_2_O_4_ and 20 mM MgCl_2_ in triplicates. After 1 h (T1) and 4 h (T2), total bacterial RNA was isolated by a hot phenol extraction protocol (see above), DNA was digested using the TURBO DNase (Ambion), purified with phenol:chlorophorm:isopropanol, and the quality was assessed using the Agilent RNA 6000 Nano Kit on the Agilent 2100 Bioanalyzer (Agilent Technologies) or Qubit (Thermo Scientific). Samples with an RIN of >8 were considered for RNA-Seq sample preparation. From the total RNA the rRNA was depleted using Ribo-Zero (Illumina). ERCC spike-ins (external RNA controls consortia Mix 1 and Mix 2) were added before rRNA depletion as a means to control for sequencing and platform performance.

Further processing included fragmentation by sonication to a median size of 200 nt, and cDNA library preparation (NEBNext Ultra II Directional RNA Library Prep Kit - Bacteria (New England Biolabs) according to the manufacturer’s instructions. Multiple oligos for Illumina sequencing (#E6440L) were obtained from New England Biolabs and used as indices that allow uniqueness in both directions. Illumina sequencing was performed with the NovaSeq 6000 sequencing system (S1 flow cell, Illumina) with 100 cycles and paired-end reads (PE50, 2 x 50 bp reads). The fluorescent images were processed into sequences and transformed to FastQ format using the Genome Analyzer Pipeline Analysis software 1.8.2 (Illumina). The sequence output was controlled for general quality features, sequencing adapter clipping, and demultiplexing using the fastq-mcf and fastq-multx tools of ea-utils [91].

### Read mapping, bioinformatics and statistics

Quality of the sequencing output was analyzed using FastQC (Babraham Bioinformatics). All sequenced libraries were mapped to the YPIII genome (NC_010465) and the pYV plasmid (accessions NC_006153) using Bowtie2 (version 2.1.0) [92] with default parameters [94]. After read mapping, SAMtools [93] was employed to filter the resulting bam files for uniquely mapped reads (both strands), which were the basis for downstream analyses. To detect ORFs and *trans*-encoded sRNAs that are differentially expressed in YPIII and the *rnc* mutant YP356, we used DESeq2 [73] for all differential expression (DE) analyses following the default analysis steps described in the package’s vignette. For each comparison, HTSeq in union count mode was used to generate raw read counts required by DESeq as the basis for DE analysis.

### Data access

The high-throughput read data is deposited at the Gene Expression Omnibus (GEO) with accession no.: GSE (https://www.ncbi.nlm.nih.gov/geo/query/acc.cgi?acc=GSE249386). The comparative transcriptome analyses are given in Datasets **S2-S4**.

### Gel electrophoresis, preparation of cell extracts, and Western blotting

For the detection of proteins of interest, *Y. pseudotuberculosis* cultures were grown under indicated growth conditions. Cell extracts of equal amounts of bacteria were prepared and separated on 12-15% SDS polyacrylamide gels [93]. The gels were either stained with Coomassie Blue or the samples were transferred onto a PVDF membrane via electro-blotting and probed with the respective antibody as described [95]. For visualization of the proteins mentioned below, generated polyclonal (Davids Biotechnology) or purchased monoclonal antibodies were used (anti-LcrF 1:1,000, gift of G. Plano; anti-CsrA 1:8000 anti-rabbit, anti-β-lactamase 1:10,000 anti-mouse, Abcam; anti-YopD 1:6,000 and anti all Yops 1:12,000 gift of A. Forsberg).

### Yop effector secretion assay

The Yop secretion assay was performed as described [96]. Bacteria were grown overnight at 25°C in LB medium, diluted 1/50 in fresh LB medium, and grown at 25°C for 2 h and then shifted to 37°C in the presence or absence of 20 mM NaOx and 20 mM MgCl_2_. Proteins in the filtered medium supernatant were precipitated with TCA and harvested by centrifugation. Precipitated proteins were washed with 95% acetone with 0.5% SDS and resuspended in equal amounts of sample buffer. Subsequently, the proteins were separated on 15% SDS polyacrylamide gels and visualized by Coomassie brilliant blue staining or Western blotting with polyclonal antibodies directed against secreted Yop proteins of *Y. pseudotuberculosis*.

### Nitrocefin-based Yop effector secretion assay

A Nitrocefin-based secretion assay was used to quantify Yop effector secretion of *Y. pseudotuberculosis* strains. For this purpose, *Y. pseudotuberculosis* strains harboring plasmid pIVO13 encoding a *yopE-blaM* fusion were grown under indicated conditions and adjusted to an OD_600_ of 0.8. Bacterial cells were pelleted and 95 µl of the supernatant were mixed with 5 μl of a 2 mmol Nitrocefin diluted in PBS and incubated for 1 h at room temperature. The absorption at 486 nm (A_486_) and 390 nm (A_390_) was measured at specific time points using the Varioskan Flash (Thermo Scientific). The specific β-lactamase activity was calculated as ratio between A_486_/A_390_ (sample) / A_486_/A_390_ (medium).

### β-galactosidase assays

**(A)** *Y. pseudotuberculosis* strains, carrying *lacZ* reporter fusions of interest were grown under indicated growth conditions in LB medium. The β-galactosidase activity of the *lacZ* fusion constructs was measured in permeabilized cells as described previously [92]. The activities were calculated as follows: β-galactosidase activity OD_415_ · 6,75 · OD_600_^-1^ ·t (min)^-1^ · Vol (ml)^-1^.

### Spotting assays

Overnight cultures were diluted 1:10 in fresh LB medium with the respective antibiotics. Bacteria were grown at 25°C for 1 hour. The OD_600_ was adjusted to 0.45 in LB medium and serial dilutions from 10^-1^-10^-7^ were prepared. 10 μl of each dilution were spotted onto LB plates and incubated at either 25°C and 37°C.

### Motility assay

A 5 µl portion of overnight culture grown at 25°C, adjusted to an OD_600_ of 0.8, was spotted onto semisolid tryptone swarm plates (1% tryptone, 0.5% yeast extract, 1% NaCl, 0.25% Difco Bacto agar). The capacity of each strain to spread was monitored after 28 hours at 25°C by measuring the diameter of the bacterial colony.

### Cell viability assay

Cell viability of bacterial cultures was measured using the BacTiter-Glo Microbial Cell Viability Assay (Promega) by detection of present ATP. Bacterial cultures were cultivated under indicated conditions and the OD_600_ adjusted to 0.8 in the growth medium. The culture was mixed 1:1 with the BacTiter-Glo substrate and luminescence was measured using the Varioskan Flash (Thermo Scientific). Cell viability was determined in duplicates by calculating the ratio of luminescence of the sample and medium.

### Scanning and transmission electron microscopy

In order to identify T3SS needles on the bacterial surface, negative staining for transmission electron microscopy and field emission scanning electron microscopy was used. Bacterial overnight cultures were diluted 1:50 in LB and grown at 25°C, 37°C, and 37°C/-Ca^2+^ for 4 h. Three independent cultures were pooled and cultures were mixed with cooled glutaraldehyde (2%) and formaldehyde (5%). For transmission electron microscopy thin carbon support films were prepared by sublimation of carbon on freshly cleaved mica. Using 300 mesh copper grids, the samples were negatively stained with 2% (w/v) aqueous uranyl acetate, according to the method of Valentine *et al.* [97], and examined in a transmission electron microscope (TEM910, Zeiss, Germany) at an acceleration voltage of 80 kV at calibrated magnifications. Images were recorded digitally with a Slow-Scan CCD-Camera (ProScan, 1024 x 1024, Scheuring, Germany) applying the ITEM-Software (Olympus Soft Imaging Solutions, Münster, Germany).

For field emission scanning electron microscopy glass coverslips were coated with a poly-L-lysine solution (Sigma, Munich, Germany), 30 µl of the samples were added, left for 10 min, washed gently with TE buffer (TRIS EDTA 0,01 M, pH 6.9) and then fixed in 2% glutaraldehyde in cacodylate buffer. After washing with TE buffer dehydration was carried out in a graded series of acetone (10, 30, 50, 70, 90, 100%) on ice for 15 min for each step. Samples were then critical-point dried with liquid CO_2_ (CPD 300, Leica, Wetzlar) and covered with a gold film by sputter coating (SCD 040, Balzers Union, Liechtenstein). For examination in a field emission scanning electron microscope (Merlin, Zeiss, Germany), an Everhart Thornley SE-detector was used with the inlens SE-detector in a 25:75 ratio at an acceleration voltage of 5 kV. Images were recorded applying the SmartSEM software version 6.06.

## Acknowledgements

We thank Dr. Martin Fenner for helpful discussions, and Tanja Krause, Bettina Elxnat, and Karin Paduch for excellent technical support.

## Supplementary Figure Legends

**Figure S1:** Influence of the loss of different RNases on growth and motility of *Y. pseudotuberculosis*.

**(A)** Overnight cultures of *Y. pseudotuberculosis* YPIII (wt) and the isogenic RNase mutants indicated on the right were diluted 1:50 in LB and growth at 25°C and 37°C was followed by measurement of OD_600_. Data represent the mean ± SD from experiments performed in triplicates. (**B**) One representative example of three independent spotting assays of serial dilutions of strains, used in **A,** on LB agar plates at 25°C is shown. (**C**) An equal amount of *Y. pseudotuberculosis* wildtype YPIII and the indicated RNase mutants was spotted onto tryptone swarm semi-soft agar plates and incubated at 25°C for 16 h. A Δ*flhD*C mutant was used as negative control (right spot).

**Figure S2:** Influence of RNase III on Yop protein secretion.

**(A)** Viability of *Y. pseudotuberculosis* strains YPIII (wt), YP356 (Δ*rnc*), and YP356 (Δ*rnc*) pIVO20 (p*rnc*^+^). Data represent the mean ± SD from four independent biological replicates (lower panel). No significant differences compared to the wildtype were determined using the Student’s t-test. (**B**) *Y. pseudotuberculosis* strains YPIII (wt), YP356 (Δ*rnc*), and YP356 (Δ*rnc*) pIVO20 (p*rnc*^+^) were grown at 37°C for 4 h; the secreted proteins in the supernatant of the cultures were precipitated with TCA and separated on SDS gels. The Yop secretion-deficient mutant YP101 (Δ*yscS*) was used as a negative control.

**Figure S3:** Influence of RNase III and PNPase on the copy number of the virulence plasmid pYV.

*Y. pseudotuberculosis* strains YPIII (wt), YP139 (Δ*pnp*), and YP356 (Δ*rnc*), were grown at 25°C, and 37°C. Total DNA of the strains grown under the different conditions was prepared and the copy number of the virulence plasmid was determined by qPCR. The relative plasmid copy number of wildtype was compared with the different mutant strains. Data represent the mean ± SD from three independent biological replicates. No significant differences were determined using Student’s t-test.

**Figure S4:** *lcrF* transcript detection.

*Y. pseudotuberculosis* (YPIII), YP356 (Δ*rnc*), and YP139 (Δ*pnp*) were grown at 37°C, and the total RNA of the strains was prepared. 10 µg of total RNA of the different strains were loaded onto a denaturing acrylamide gel to allow the detection of smaller degradation products. The *lcrF* transcripts were identified by Northern blot using a probe covering the *lcrF* coding sequence. The blot represents one of two biological replicates. The 5 S rRNA was used as loading control.

**Figure S5:** Analysis of the expression of the Yops and LcrF at different time points after induction of the T3SS.

*Y. pseudotuberculosis* (YPIII) was grown over day at 25°C to exponential phase and was then shifted to 37°C in the presence (37°C) or absence of Ca^2+^ (-Ca^2+^). Whole-cell extracts were prepared and the amounts of Yops (**A**) and LcrF (**B**) were analyzed by Western blotting using all-Yop and LcrF polyclonal antisera. H-NS was used as loading control. The Δ*lcrF* mutant was used as a negative control.

**Figure S6:** RNA-Seq quality control. (**A**) Two-dimensional biplot and (**B**) three-dimensional principal component analysis of mean-centered and scaled rlog-transformed read count values of the *Y. pseudotuberculosis* wildtype strain YPIII and the isogenic Δ*rnc* mutant grown over day at 25°C to exponential phase (T0) and then shifted to 37°C in the presence or absence of Ca^2+^ for 1 h (T1) or 4 h (T2).

